# Macrophage-specific lipid nanoparticle therapy blocks the lung’s mechanosensitive immunity due to macrophage-epithelial interactions

**DOI:** 10.1101/2023.05.24.541735

**Authors:** Liberty Mthunzi, Mohammad N Islam, Galina A Gusarova, Sunita Bhattacharya, Brian Karolewski, Jahar Bhattacharya

**Affiliations:** Lung Biology Laboratory, Division of Pulmonary, Allergy, and Critical Care Medicine, Department of Medicine, Vagelos College of Physicians and Surgeons, Columbia University, New York, NY 10032, USA; Institute of Comparative Medicine, Columbia University Irving Medical Center, Columbia University, New York, NY 10032, USA; Department of Physiology and Cellular Biophysics, Columbia University, New York, NY 10032, USA

**Author notes:** Corresponding author: Jahar Bhattacharya, 630 West 168^th^ St, BB8-812. Co-first authors (Liberty Mthunzi, Mohammad N Islam).

## Abstract

The lung’s mechanosensitive immune response, which occurs when pulmonary alveoli are overstretched, is a major impediment to ventilation therapy for hypoxemic respiratory failure. The cause is not known. We tested the hypothesis that alveolar stretch causes stretch of alveolar macrophages (AMs), leading to the immune response. In lungs viewed by optical imaging, sessile AMs expressed gap junctional protein connexin-43 (Cx43), and they communicated with the alveolar epithelium through gap junctions. Alveolar hyperinflation increased Ca^2+^ in the AMs but did not stretch the AMs. The Ca^2+^ response, and concomitant TNFα secretion by AMs were blocked in mice with AM-specific deletion of Cx43. The AM responses, as also lung injury due to mechanical ventilation at high tidal volume, were inhibited by AM-specific delivery of lipid nanoparticles containing Xestospongin C, which blocked the induced Ca^2+^ increases. We conclude, Cx43- and Ca^2+^-dependent AM-epithelial interactions determine the lung’s mechanosensitive immunity, providing a basis for therapy for ventilator-induced lung injury.

## INTRODUCTION

Mechanical ventilation, critical therapy in hypoxemic acute lung injury (ALI), which leads to ARDS (1), supports blood oxygenation and lung recovery. However, mechanical ventilation is itself proinflammatory in that it can cause a mechanosensitive immune response that recruits inflammatory cells to the lung’s air spaces, resulting in injury to the air-blood barrier (2, 3). This mechanical injury reverses the protective effects of ventilator therapy and contributes to morbidity and mortality in ARDS (2, 3). It is proposed that mechanical injury results from alveolar overexpansion during mechanical ventilation (3, 4). However, the mechanistic link between alveolar overexpansion and the immune response remains poorly defined, hampering development of precision therapy for patients selected for mechanical ventilation.

Macrophages are implicated in the lung’s mechano-immune response. However, this view is based on *in vitro* evidence that macrophage stretch induces Ca^2+^ influx, potentially a proinflammatory response, across mechanosensitive ion channels (5, 6). In heart, *in vitro* evidence for mechanosensitive mechanisms implicates focal adhesion dependent cell-tissue contacts that transmit tissue stretch to macrophages (7). Alveolar expansion might similarly transmit stretch from epithelium to alveolus-adherent, sessile alveolar macrophages (AMs) (8), activating proinflammatory signaling. However, supportive evidence for this possibility is lacking.

Understanding of the AM stretch hypothesis *in situ* is hampered by two major knowledge gaps. These are first, the extent to which the non-uniformity of alveolar expansion, as revealed by real-time confocal microscopy (RCM) of mouse lungs (9), impacts stretch transmission to sessile AMs remains undefined. Second, the extent to which expansion is also non-uniform in human alveoli remains unconfirmed. Hence, from the standpoint of mouse models of alveolar overexpansion and their translational relevance, the role of macrophage stretch as an activator of the lung’s mechano-immunity remains unclear.

Here, we tested the AM stretch hypothesis by RCM of live lungs to assess hyperinflation-induced distension of alveolar segments that contained or lacked sessile AMs. For translational relevance, we determined alveolar expansion patterns in live, transplant-rejected human lungs as we described previously (10). Since human lungs are relatively in short supply, we also determined responses in pristine pig lungs. Being comparably sized with human lungs, pig lungs provide a model for translational studies of mechanical ventilation (11, 12). An associated goal was to define the extent to which hyperinflation activated calcium (Ca^2+^) influx in the AMs, thereby priming the proinflammatory response. We report below our unexpected finding that the lung’s mechano-immunity is determined by a Ca^2+^-dependent epithelial-macrophage crosstalk that licenses the immune response to a subset of sessile AMs. We also report a therapeutic approach for suppressing this signaling.

## RESULTS

### Alveolar expansion does not change cell shape in sessile AMs

To determine the effects of alveolar expansion on cell shape of sessile AMs, we viewed alveoli in mouse, pig and human lungs by RCM. Alveolar microinfusions of calcein-AM (calcein) induced alveolar epithelial fluorescence, marking the alveolar perimeter (8) (Fig. 1a-c and sFig. 1a). Calcein also marked sessile AMs, which were lumen-facing, alveolus-adherent cells. AMs were further identified by specific markers, namely SiglecF and CD11c for mouse, Siglec1 for human and CD203a for pig (sFig. 1a) (8, 13, 14). As we reported previously (8), the AMs remained stationary.

**Figure 1.**
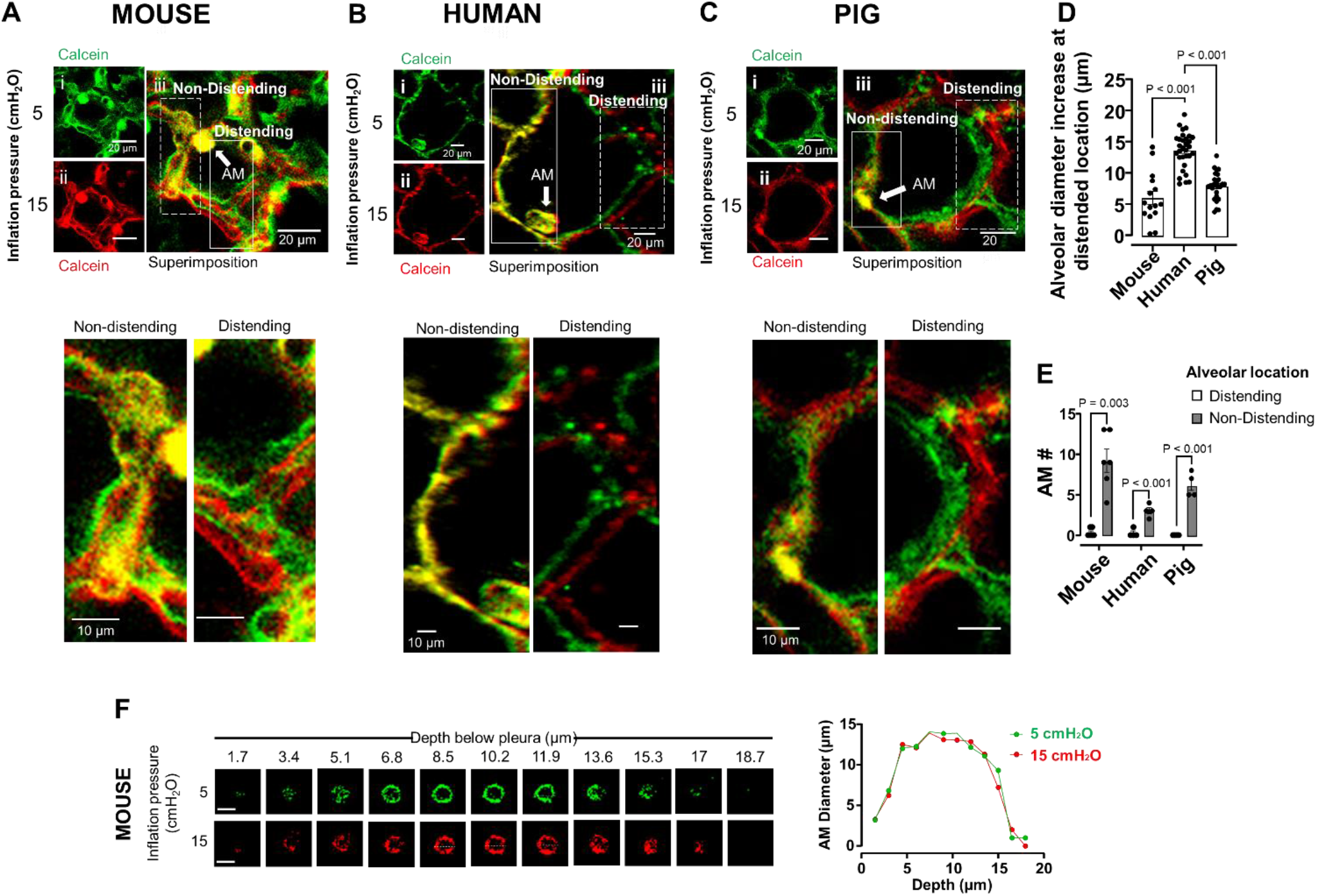
Pseudocolor superimposition by confocal microscopy reveals AM shape protection during mouse, human and pig lung hyperinflation **a-c** Representative confocal images of mouse (a) human (b) and pig (c) alveoli are shown at low *(i-ii)* and high magnification (*iii*). Images were obtained at the indicated inflation pressures. The low magnification images are pseudocolored differently for the different inflation pressures. The merge image (*iii*) was obtained by superimposing the green and red images. The superimposition reveals yellow pseudocolor at a non-distending alveolar segment (*solid rectangle*), but separation of red and green at the distending segment (*dashed rectangle*). A sessile alveolar macrophage (*AM*) is located at the corner of the non-distending alveolar segment (*arrow*). High magnification images of rectangles are below. **d** Bars show hyperinflation-induced alveolar diameter increase at distending location in mouse, human and pig lungs. n = 15-40 alveoli from 4-6 lungs per group. Mean±sem. **e** Bars show quantification of AMs at distending and non-distending locations of alveoli mouse, human and pig lungs. Each dot shows number of AMs data from one lung. n = 4-6 lungs per group. Mean±sem. **f** Confocal z stack images show the mouse AM perimeter (SiglecF staining) at 5 (green) and 15 (red) cmH_2_O inflation pressure at different depths. The graph plots AM diameter (dashed line) at different depths below the pleura. Replicated 6 times in 6 lungs. Scale bar = 10 µm.

To quantify alveolar shape changes, we applied our reported approach in which we image an alveolus at low and high inflation pressures, using a different pseudocolor for each image (9). Superimposition of the images reveals congruence, or separation of the pseudocolors at respectively, non-distended or distended alveolar segments. We increased inflation pressure from 5 to 15 cmH_2_O, a procedure that increases lung volume to about 80% of total lung capacity (9). We avoided very high airway pressures, such as those applied in previous studies (15), to eliminate the risk of disruptive alveolar injury and bleeding. The pressure increase caused uneven alveolar expansion not only in mouse lungs, as we reported (9), but also in human and pig lungs (Fig. 1a-c). At the distended locations the alveolar diameter increased by several micrometers (Fig. 1d).

Our new finding was that sessile AMs were almost exclusively located at alveolar regions that did not distend after hyperinflation (Fig. 1e). AM diameter was smaller in mouse (12±2 μm) and pig (11±1 μm) than in human (21±2 μm) (sFig. 1b). Despite these morphological differences, superimposed pseudocolors obtained at different inflation pressures were completely merged for mouse, human and pig AMs (sFig. 1c-e), suggesting that at a single optical level, hyperinflation did not change AM diameter. We carried out optical imaging at different optical levels down the depth axis of each AM (sFig. 1f), then at each optical level we quantified AM diameter which we plotted as a function of depth. We expected that to the extent that hyperinflation changed AM shape, the plots for baseline and hyperinflation would be non-congruent. However, the plots were completely congruent for mouse (Fig. 1f), human and pig AMs (sFig. 1g, h), indicating that hyperinflation did not cause AM shape changes.

Taken together, these findings indicate that in mouse, human and pig lungs, since AMs were located at non-distending alveolar segments, epithelium-AM stretch transmission was absent. Thus, alveolar overexpansion did not cause AM stretch.

### Hyperinflation mobilizes Ca^2+^ in a subset of sessile AMs

To determine hyperinflation-induced immune mechanisms alternative to AM stretch, we quantified AM Ca^2+^. Following steady levels at baseline, a 15-second hyperinflation due to airway pressure increase from 5 to 15 cmH_2_O, induced prolonged Ca^2+^ increases (Fig. 2a, b). The subsequent return of Ca^2+^ to baseline ruled out the possibility that the response was injury induced. Ca^2+^ increases did not occur when we increased airway pressure from 5 to 10 cmH2O (sFig. 2), indicating that the Ca^2+^ response was not progressive.

**Figure 2.**
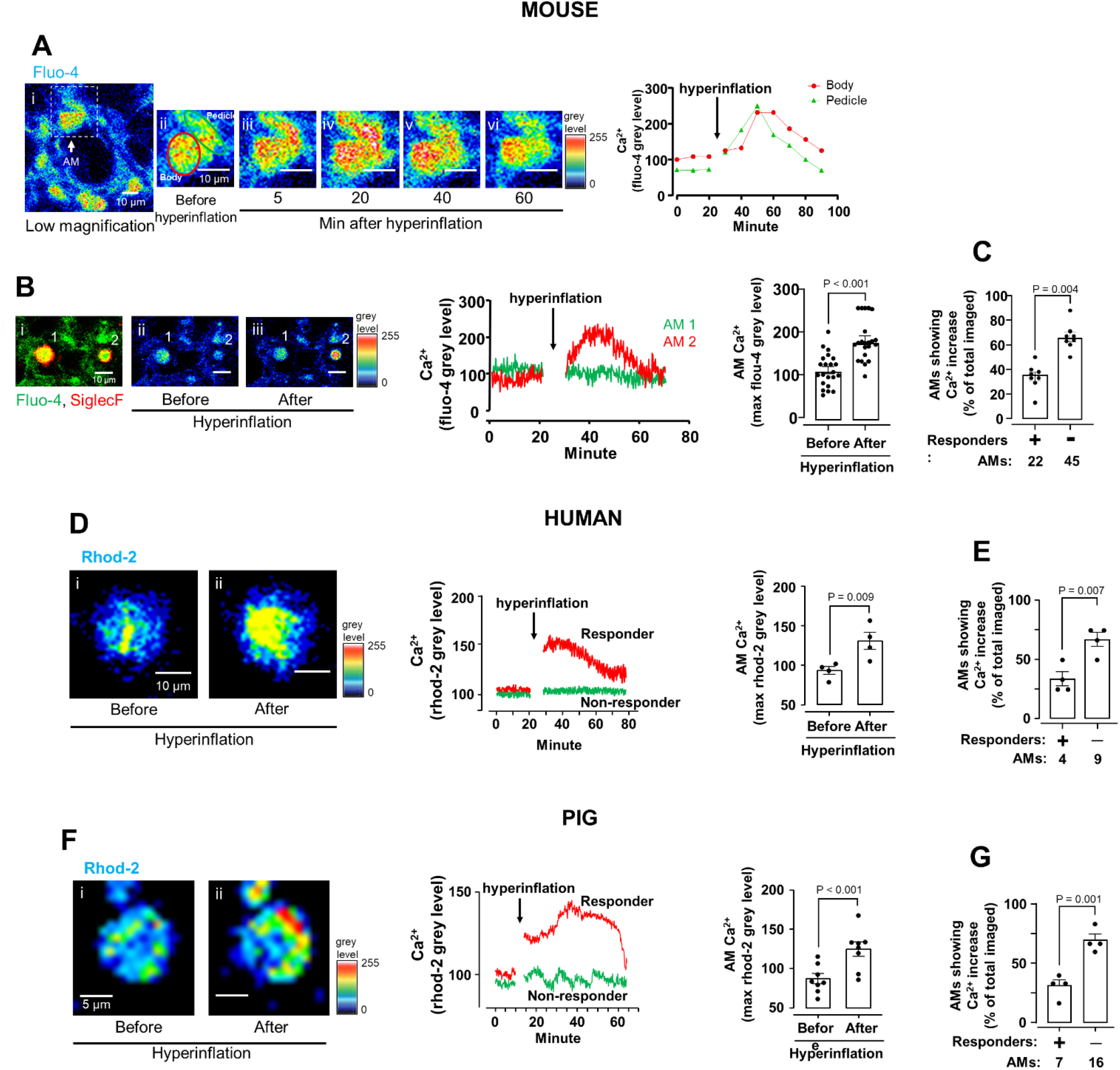
Hyperinflation causes calcium mobilization in mouse, human and pig sessile AMs **a** Confocal images of mouse alveolar epithelium loaded with the fluorescent cytosolic calcium indicator, fluo-4 show cytosolic Ca^2+^ fluorescence in a single AM at low (*rectangle in i*) and high (*ii-vi*) magnification before (*i-ii*) and after hyperinflation (*iii-vi*). Pseudocolors represent fluorescence expressed as grey levels as defined in the symbol key. Tracings show Ca^2+^ in cell body and pedicle (red and green circles in *ii*). **b** Images from adjoining alveoli show AM that responded with hyperinflation-induced Ca^2+^ increase (AM #2) versus an AM that did not respond (AMs #1). Tracings show corresponding time courses. Bars show paired comparisons of maximum Ca^2+^ before and after hyperinflation in AMs that responded to hyperinflation. **c** Bars show paired comparisons for responders (+) and non-responders (-) from 8 lungs. Mean±sem. **d** Confocal images of human AMs loaded with the fluorescent mitochondrial calcium indicator, rhod-2 show mitochondrial Ca^2+^ fluorescence in a single AM at high magnification before (*i*) and after hyperinflation (*ii*). Pseudocolors represent fluorescence expressed as grey levels as defined in the symbol key. Tracings show mitochondrial Ca^2+^ in AMs. Bars show paired comparisons of maximum Ca^2+^ before and after hyperinflation in AMs that responded to hyperinflation. **e** Bars show paired comparisons for responders (+) and non-responders (-) from human 4 lungs. Mean±sem. **f** Confocal images of pig AMs loaded with the fluorescent mitochondrial calcium indicator, rhod- 2 show mitochondrial Ca^2+^ fluorescence in a single AM at high magnification before (*i*) and after hyperinflation (*ii*). Pseudocolors represent fluorescence expressed as grey levels as defined in the symbol key. Tracings show mitochondrial Ca^2+^ in AMs. Bars show paired comparisons of maximum Ca^2+^ before and after hyperinflation in AMs that responded to hyperinflation. **g** Bars show paired comparisons for responders (+) and non-responders (-) from 4 pig lungs. Mean±sem.

Although hyperinflation expanded all mouse alveoli, Ca^2+^ increases occurred in 35% of the AMs (Fig. 2c). As exemplified, AMs that lacked Ca^2+^ responses were frequently next-alveolus neighbors of AMs that responded with Ca^2+^ increases (Fig. 2b). We conclude, although hyperinflation caused epithelial stretch in all alveoli, the induced Ca^2+^ increases occurred in a subset of sessile AMs.

Human and pig alveoli were not edematous or injured and they inflated well as delineated by epithelial fluorescence of calcein which denoted cellular viability (Fig. 1b, c). Nevertheless, in these alveoli, cytosolic Ca^2+^ detection was not possible as Fluo-4 was not internalized by cells. In alveolar type 2 cells, mitochondrial buffering masks cytosolic Ca^2+^ increases due to lung hyperinflation (16). The buffering increases the mitochondrial Ca^2+^, which is therefore a surrogate for the cytosolic response. Since the mitochondrial Ca^2+^ indicator, Rhod-2 was internalized by alveolar cells of human and pig lungs, we opted for mitochondrial Ca^2+^ detection in these lungs.

Our findings indicate that a third of all AMs tested in mouse, human and pig lungs responded with substantial Ca^2+^ increases to hyperinflation (Fig. 2b-g). We note that the similarity of these responses suggests translational relevance of the mouse findings. However, this interpretation needs to be made with caution and further studies designed to improve translational understanding.

### Hyperinflation causes store release of Ca^2+^ in sessile AMs

To determine mechanisms underlying the hyperinflation-induced Ca^2+^ increase, we affirmed that when a single hyperinflation was followed by a second hyperinflation, each challenge induced similar Ca^2+^ transients (Fig. 3a). However, given by alveolar microinfusion after the first hyperinflation, Xestospongin C (XeC) inhibited the Ca^2+^ increase due to the second hyperinflation. Since XeC blocks store release (17), we interpret that hyperinflation-induced Ca^2+^ increases resulted from endosomal Ca^2+^ release.

**Figure 3.**
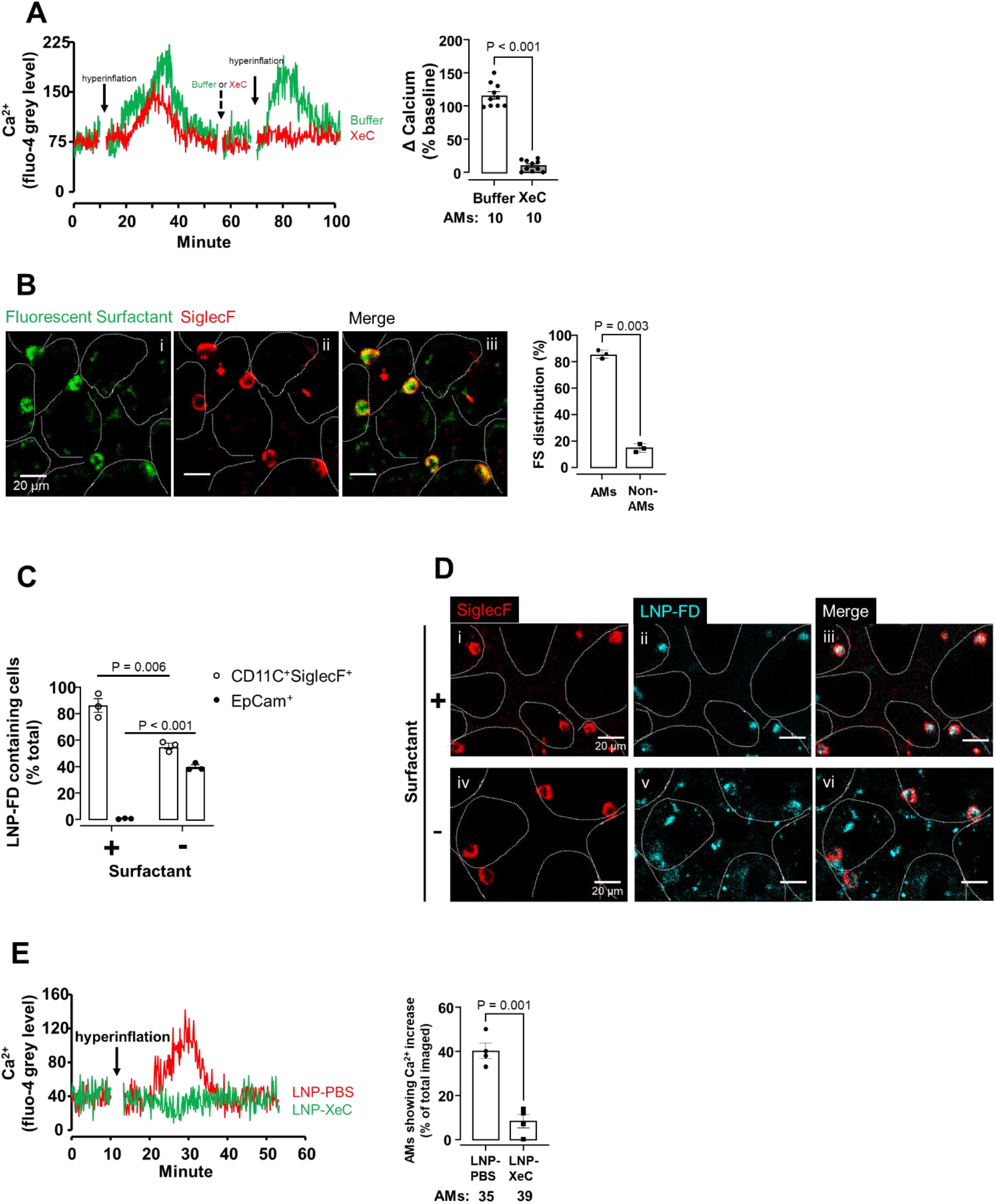
Targeted delivery of Xestospongin C to AMs inhibits calcium mobilization **a** Tracings in different colors show Ca^2+^ responses in different sessile AMs following sequential hyperinflation challenges as shown (solid arrows). Alveolar microinfusions of buffer alone (green tracing), or buffer containing xestospongin C (*XeC*) (red tracing) were given as shown (dashed arrow) in different lungs. Bars show the group data following second hyperinflation. Mean±sem, 10 AMs from 3 lungs for *Buffer* group and 10 AMs from 4 lungs for *XeC* group. **b** Confocal images show uptake of fluorescent surfactant (FM1-43 labelled) (*i*) in AMs (*ii-iii*) 2 hours after intranasal instillation. Bars show quantification of uptake from 3 lungs. Mean±sem. Dotted lines trace alveolar perimeters. *FS*, fluorescent surfactant. **c** Bars show flow cytometry quantification of rhodamine b dextran 70 kDa LNPs (LNP-FD) uptake in alveolar macrophages (CD11c+SiglecF+) and alveolar epithelium (EpCam+) from lung cell suspension 2 hours after instillation. n = 3 lungs. *LNP-FD*, lipid nanoparticle-encapsulated fluorescent dextran. Mean±sem. **d** Confocal images show AM (SiglecF+) uptake of lipid nanoparticle-encapsulated rhodamine b labelled dextran 70 kDa (LNP-FD) 2 hours after intranasal instillation in presence (+, *i-iii*) or absence of surfactant (-, *iv-vi*). **e** Tracings show Ca^2+^ response in AMs of mice intranasally instilled with lipid nanoparticles encapsulating PBS (*LNP-PBS*) or Xestospongin C (LNP-*XeC*) 2 hours prior to lung imaging. Bars show quantification of the % of AMs responding with Ca^2+^ increase post hyperinflation in AMs of mice intranasally instilled with *LNP-PBS* or *LNP-XeC* 2 hours prior to lung imaging. Mean±sem, n = 4 lungs per group.

Since alveolar microinfusion does not ensure cell-specific delivery, we developed a surfactant-based strategy (18) for XeC delivery to AM. Intranasally instilled surfactant, which was fluorescently labelled with FM1-43, was engulfed primarily by AMs (Fig. 3b), indicating that exogenous surfactant potentially provided a vehicle for AM-specific drug targeting. We prepared lipid nanoparticles that encapsulated fluorescent dextran (LNP-FD), then gave the LNP-FD with surfactant. For control, we gave LNP-FD without surfactant.

Our flow cytometry findings indicate that AMs comprised 90% of cells that internalized surfactant-associated LNP-FD (Fig. 3c). Importantly, LNP-FD was absent in EpCAM-positive cells (Fig. 3c), indicating that the alveolar epithelium did not internalize the nanoparticles. It is possible that about 10% of the uptake occurred in other cells, such as neutrophils. However, we did not detect LNP-FD uptake in spleen, heart and liver (sFig. 3b), indicating that surfactant-associated LNP-FD were not taken up in systemic organs. By contrast, given without surfactant, LNP-FD were extensively taken up in the alveolar epithelium (Fig. 3c).

These findings were affirmed by RCM data which showed that surfactant-associated LNP-FD localized to AMs, while given without surfactant LNP-FD were extensively taken up by other cells (Fig. 3d). Thus, inclusion of surfactant in the instillation was necessary to specifically target LNP delivery to AMs and importantly, to avoid non-specific delivery to other cell types.

The success of this macrophage-targeted delivery system motivated us to suspend LNPs containing XeC in surfactant (LNP-XeC). RCM studies in lungs of these mice indicated marked inhibition of hyperinflation-induced Ca^2+^ increase in sessile AMs (Fig. 3e). As we showed previously (19), hyperinflation increased Ca^2+^ oscillations in the alveolar epithelium (sFig. 4a).

**Figure 4.**
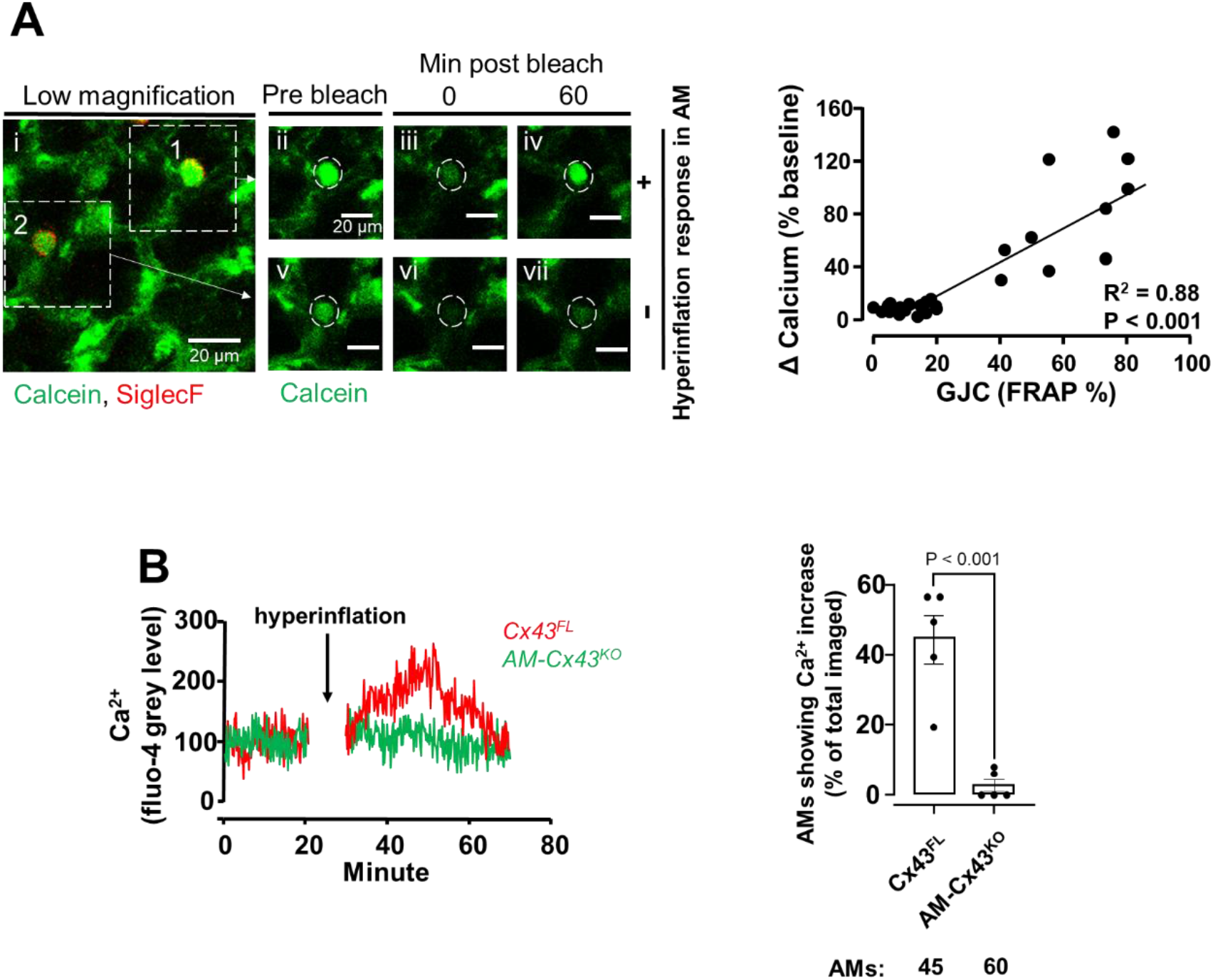
Role for connexin-43 in AM calcium mobilization **a** Confocal image (*i*) of alveolar epithelium and SiglecF positive sessile AMs loaded with the cytosol localizing dye, calcein. Dashed squares enclose two sessile AMs subjected to photobleaching. High magnification images (*ii-vii*) show calcein fluorescence in a hyperinflation responsive AM (#*1, ii-iv*) and non-responsive AM (#*2, v-vii*) pre, and post bleach at indicated time points. Plot shows regression of increase in cytosolic Ca^2+^ in AMs after hyperinflation (*Δ calcium % baseline*) against efficiency of gap junctional communication (*GJC*) quantified in terms of fluorescence recovery after photobleaching (*FRAP %*). Data are for 27 sessile AMs from 4 lungs. **b** Tracings in different colors show Ca^2+^ responses in sessile AMs from indicated mice. Bars show % of sessile AMs per field of view that responded to hyperinflation by increasing Ca^2+^ in CD11cCre-Cx43^fl/fl^ mice (*AM-Cx43^KO^*) and littermate controls (*Cx43^FL^*). Data show mean±sem. n = 5 lungs per group. Data analyzed by non-paired t-test.

However, these Ca^2+^ oscillations were not blocked by LNP-XeC (sFig. 4b), indicating that the surfactant-associated delivery avoided epithelial effects. Taking these findings together, we conclude that LNP-XeC were AM targeted and that the AM Ca^2+^ response to hyperinflation was initiated by Ca^2+^ release from endoplasmic stores.

### Connexin-43 determines Ca^2+^ mobilization in sessile AMs

In several experiments, as for in the example shown (Fig. 2a), an AM pedicle was evident that appeared to anchor the AM on the epithelium. In these instances, the hyperinflation-induced Ca^2+^ increase occurred first in the pedicle before proceeding to the cell body of the AM (Fig. 2a). These findings suggested that hyperinflation may have induced a Ca^2+^ wave that followed a trajectory from the pedicle to the cell body.

Sessile AMs establish gap junctional communication (GJC) with the alveolar epithelium through connexin-43 (Cx43)-containing gap junctions (8). Hence, we assessed efficacy of GJC by quantifying fluorescence recovery after photobleaching (FRAP) (8). Our findings indicate that the hyperinflation-induced Ca^2+^ response correlated positively with GJC efficacy between AMs and the epithelium. Ca^2+^ increases were absent in AMs with reduced GJC (Fig. 4a). In transgenic mice lacking Cx43 in sessile AMs (8) (AM-Cx43^KO^), which therefore lacked GJC (sFig. 5), hyperinflation-induced Ca^2+^ increases were absent as compared with floxed littermate controls (Cx43^FL^) (Fig. 4b). Together, these findings indicated that AM-epithelium communication by Cx43 dependent GJC sufficiently accounted for the hyperinflation-induced Ca^2+^ increases in sessile AMs, further ruling out macrophage stretch as the underlying mechanism.

**Figure 5.**
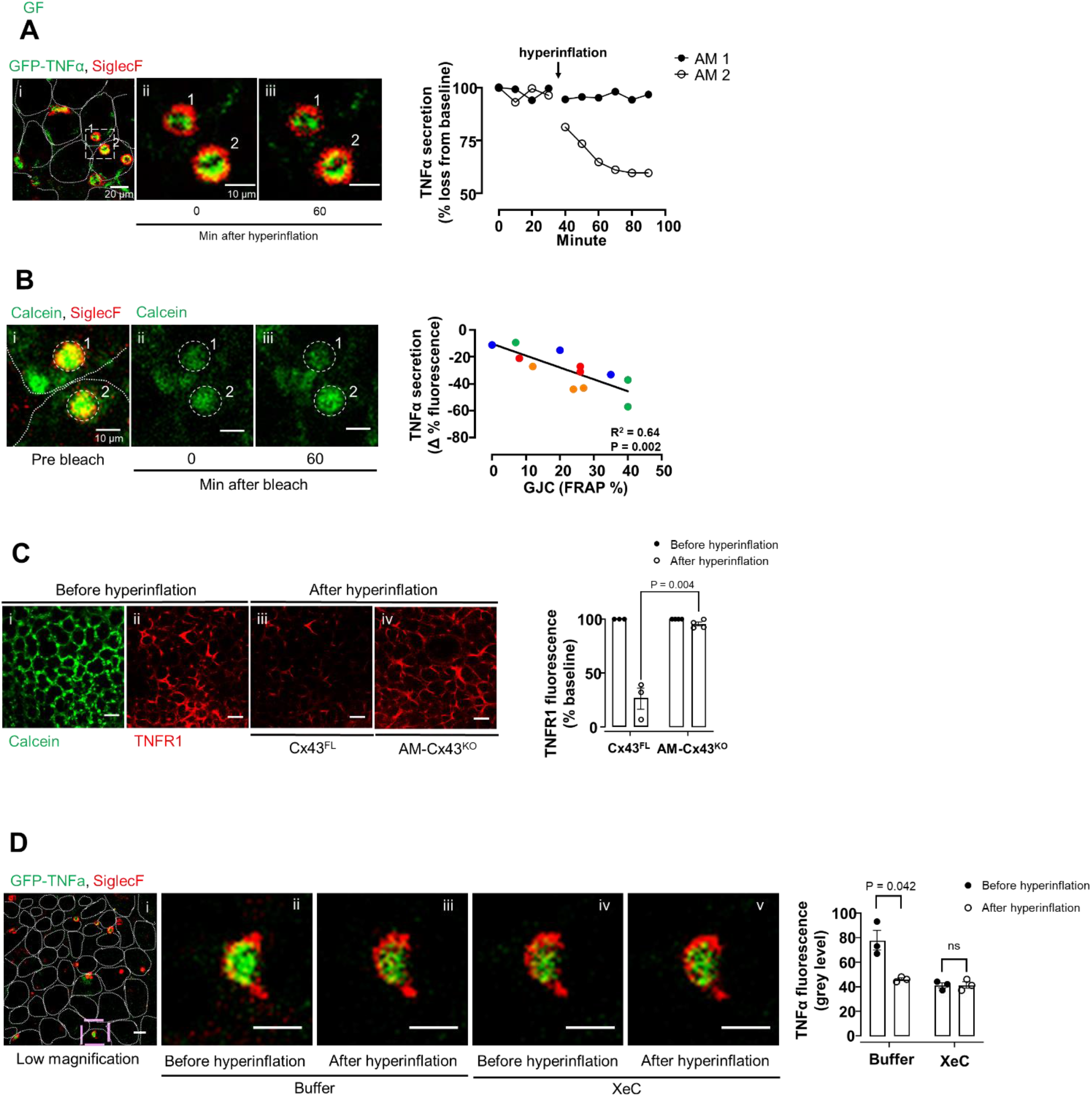
Hyperinflation induces TNFα secretion in sessile AMs **a** Confocal image (*i*) show overlay of the indicated fluorescence labels on sessile AMs. Dotted lines indicate alveoli in which sessile AMs are located. Selected AMs (rectangle), imaged at high magnification (*ii, iii*), show TNFα fluorescence before and after hyperinflation. The tracings depict fluorescence quantification at the indicated times. **b** Confocal image (*i*) shows overlay of the indicated fluorescence labels on sessile AMs from same imaging field as **a**. Dotted lines indicate alveolar epithelial margins. Confocal images (*ii-iii*), show calcein fluorescence alone after laser bleach at indicated times. Dashed circles indicate laser bleached regions. Regression plot shows data for individual AMs from 3 lungs. Same color indicates AMs from an individual lung. TNFα secretion is quantified as percent fluorescence decrease from baseline. *GJC*, gap junction communication; *FRAP %*, fluorescence recovery after photobleaching expressed as difference from initial fluorescence, quantified 60 min after bleaching. Line is drawn by linear regression analysis. **c** Confocal images of epithelial and TNFR1 fluorescence in alveoli microinfused with calcein (*i*) and anti-TNFR1 mAb (*ii-iv*) in littermate control mice (*Cx43^FL^*) and in mice with AM-specific Cx43 knockout (*AM-Cx43^KO^*) as indicated. Scale bars = 10 μm. Bars show the group data. Mean±sem. n = 3 and 4 lungs respectively, for littermate control and KO groups. Data analyzed by unpaired t-test followed by Bonferroni correction. **d** Confocal image (*i*) shows overlay of the indicated fluorescence labels on sessile AMs. Selected AM (purple rectangle in *i*), imaged at high magnification (*ii-v*), show TNFα fluorescence before and after hyperinflation in absence (*Buffer*) or presence (*XeC*) of Xestospongin. Bars show quantification of TNFα secretion. n = 3 lungs. Mean±sem.

### Mechanosensitive TNFα secretion occurs in sessile AMs

A transient Ca^2+^ increase induces vesicular exocytosis, hence secretion (20). Hence, we next asked whether the hyperinflation-induced Ca^2+^ increase in sessile AMs was of sufficient duration and magnitude as to activate secretion of TNFα, the critical initiator of proinflammatory responses. In macrophages transfected *in vitro* with a plasmid expressing GFP-TNFα (21), LPS or cytokine exposure initiates TNFα secretion with a delay of hours during which TNFα-containing vesicles dock on the cell membrane (21). Although hyperinflation increases TNFα levels in the bronchoalveolar lavage (22, 23), it is not clear whether AM interactions with neighboring cells determine the secretion.

To evaluate this question, we intranasally instilled lipid nanoparticles encapsulating GFP-TNFα plasmid in surfactant to transfect sessile AMs *in vivo*. After 48h, a majority of sessile AMs expressed TNFα fluorescence with a polar distribution in that the fluorescence was dominantly expressed at the lumen-facing cytosol (Fig. 5a and sFig. 6). Hyperinflation rapidly decreased TNFα fluorescence (Fig. 5a), indicating that the cytokine was secreted. However, the secretion response was absent in several AMs. FRAP analyses revealed a positive correlation between the extent of the secretory response and GJC efficacy (Fig. 5b). Further, shedding of alveolar TNFα receptor-1 (TNFR1), a marker of TNFα ligation (24), was blocked in AM-Cx43^KO^ mice (Fig. 5c). To determine the role of AM Ca^2+^ in the secretion response, we gave alveolar microinfusion of buffer or XeC prior to hyperinflation. XeC, but not buffer blocked hyperinflation-induced TNFa secretion in AMs (Fig. 5d), indicating that the secretory response resulted from Ca^2+^ release from endoplasmic stores.

**Figure 6.**
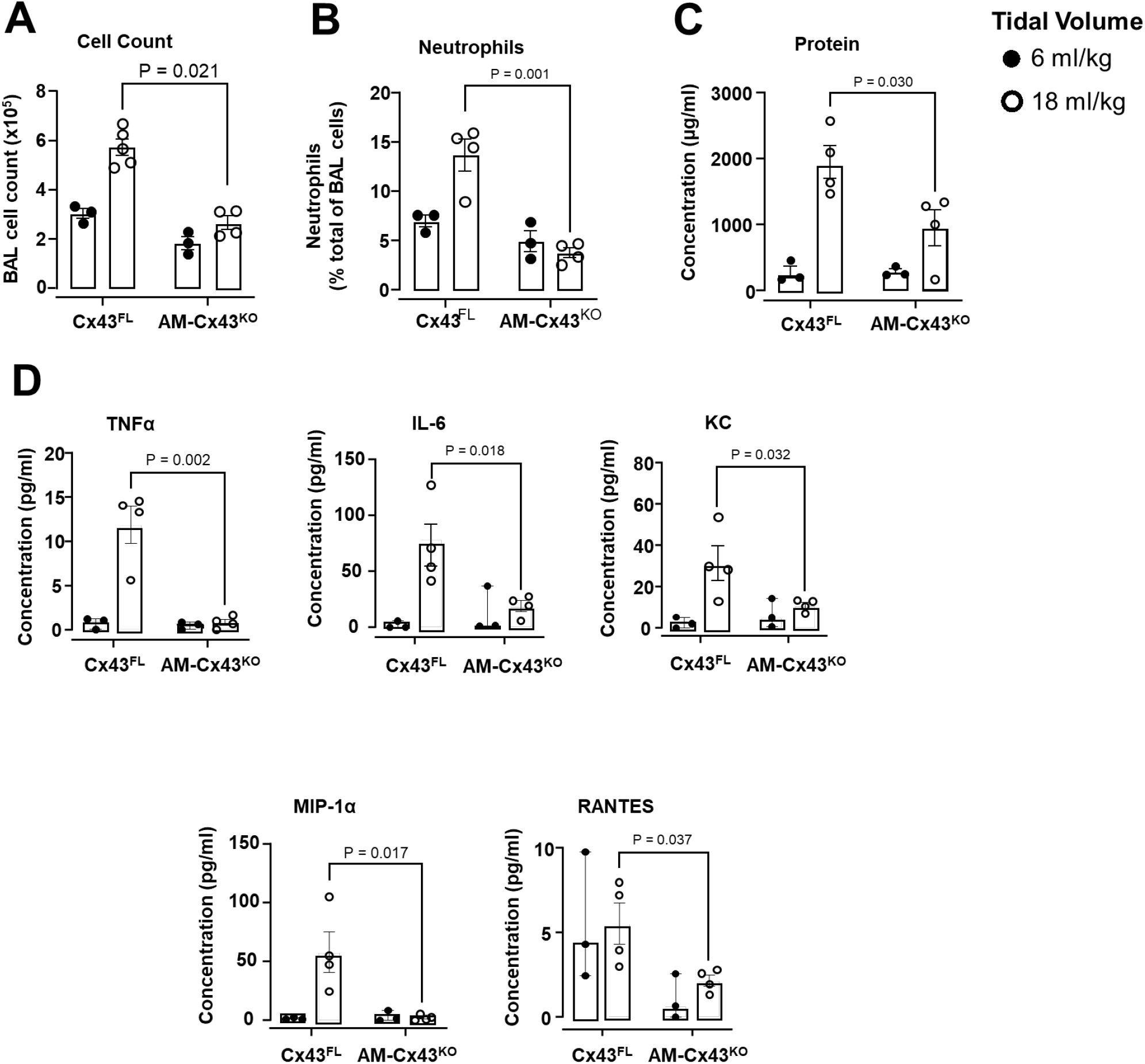
Sessile AM connexin-43 deletion protects against high tidal volume ventilation-induced lung injury **a-d** Bars show quantifications in the bronchoalveolar lavage (BAL) for total cells (a) neutrophil content (b), protein concentration (c), cytokine and chemokine levels (d) in mice 2 hours after mechanical ventilation at indicated tidal volumes. *Cx43^FL^,* littermate control. *AM-Cx43^KO^*, CD11cCre-Cx43^fl/fl^. n = 3 (*Cx43^FL^*), 4-5 (*AM-Cx43^KO^*) lungs per group. Data analyzed by unpaired t-test followed by Bonferroni correction for multiple comparisons where necessary.

Taking our findings together, we interpret that inhibition of GJC in the knockout mice blocked the Ca^2+^ increase, hence release of TNFα that ligated TNFR1. Thus, GJC between AMs and the epithelium was critical for the hyperinflation-induced TNFα secretion.

### Lung injury induced by high tidal volume ventilation results from GJC between epithelium and AMs

Mice mechanically ventilated at high tidal volume (HTV) develop lung inflammation and alveolar epithelial injury (25). To determine underlying mechanisms, we ventilated anesthetized mice at HTV to induce lung hyperinflation with each breath for prolonged durations of ventilation. Our goal was to match the lung hyperinflation due to HTV with that for the single hyperinflation applied in excised lungs. Hence, we mechanically ventilated mice at HTV (18 ml/kg), or low tidal volume (LTV) (6 ml/kg). The end-inspiration airway pressure for HTV was higher than LTV by about 10 cmH2O, similar to the pressure increase we induced in the single hyperinflation studies. Further, since mouse TLC is approximately 1 ml (26), the HTV tidal volume (∼540 ul), added to mouse FRC of ∼250 μl (26), accounts for lung expansion to about 80% TLC in a 30 g mouse. Thus, the lung expansion in HTV matched that for the single hyperinflation studies. Also similar to the single hyperinflation response, after 30 min of HTV ventilation in excised lungs, AM Ca^2+^ increased markedly above baseline, then gradually decreased in ∼1 hour (sFig. 7), indicating that the Ca^2+^ responses in the two models were similar.

**Figure 7.**
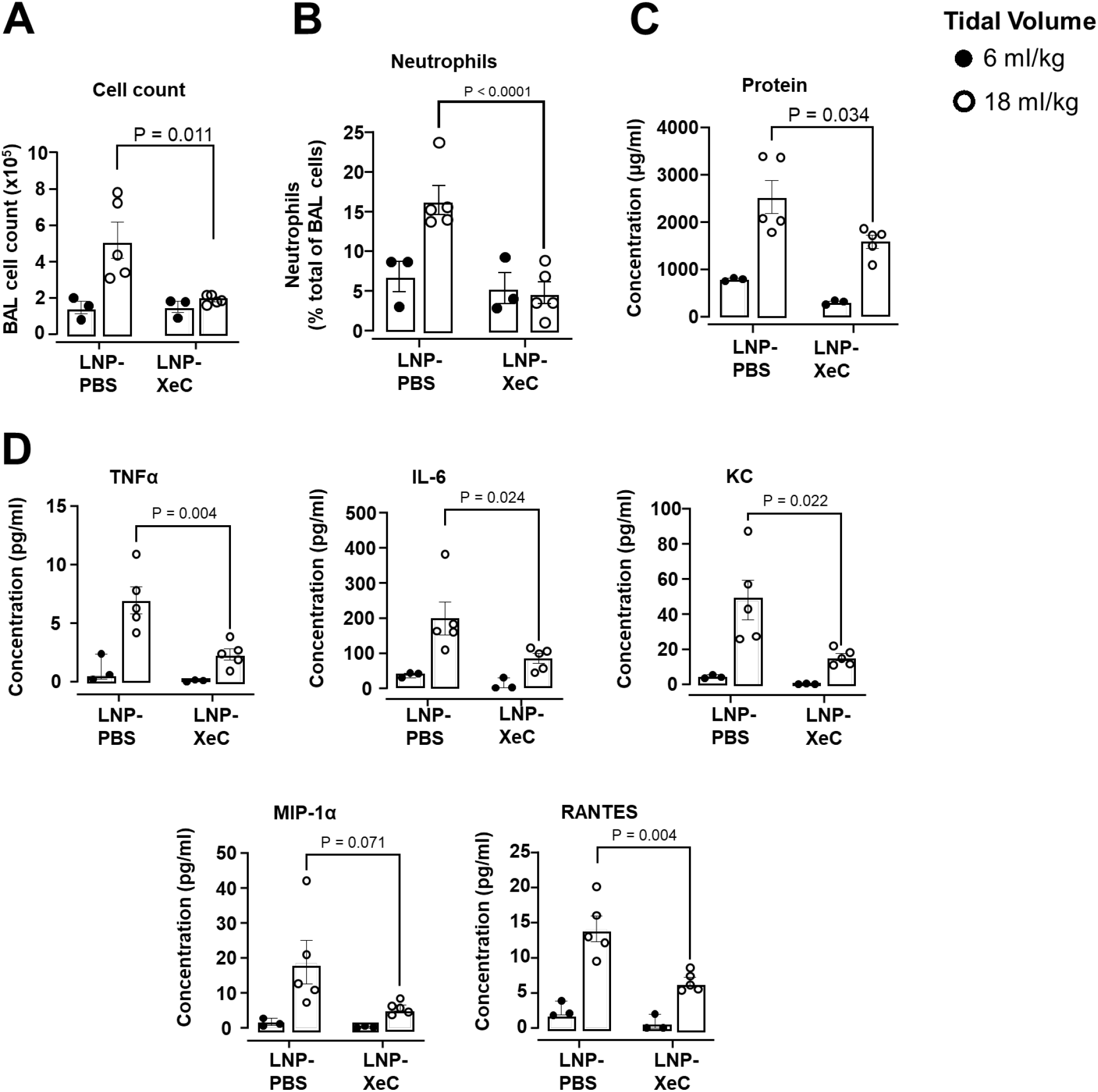
LNP-XeC pre-treatment increases protects against high tidal volume ventilation-induced lung injury **a**-**d** Bars show quantifications in the BAL for total cells (a), neutrophil content (b), protein (c), and the indicated cytokines and chemokines (d) 2 hours after mechanical ventilation at indicated tidal volumes. *LNP-XeC*, mice treated with lipid nanoparticles (LNPs) encapsulating XeC, LNP-*PBS*, mice treated with LNPs encapsulating PBS. n = 3-5 (LNP-*PBS*) and 3-5 (LNP- *XeC*) lungs per group. LNPs were given in a surfactant containing solution. Data for c-e analyzed by unpaired t-test followed by Bonferroni correction for multiple comparisons where necessary.

In Cx43^FL^ mice, two hours of mechanical ventilation at HTV induced major lung inflammation as indicated by increased content of total cells and neutrophils in the bronchoalveolar lavage (BAL) (Fig. 6a, b). In addition, BAL levels of the proinflammatory factors, tumor necrosis factor alpha (TNFα), IL-6, Keratinocyte Chemoattractant (KC), Macrophage Inflammatory Protein-1 alpha (MIP-1α /CCL3) and RANTES (CCL5) increased two- to three-fold above baseline (Fig. 6d). Since 2 hours of HTV does not cause pulmonary edema, as assessed by the extravascular lung water content (25), we quantified alveolar barrier permeability in terms of the BAL protein content, which increased more than two times (Fig. 6c). These injury responses were markedly inhibited in AM-Cx43^KO^ mice. We conclude, Cx43 expression in sessile AMs critically determined HTV-induced lung inflammation and injury.

Since LNP-XeC instillation in surfactant blocked AM Ca^2+^ increases, we gave the instillation 2 hours prior to HTV. In a separate group, we gave HTV for 1 h, instilled LNP-XeC in surfactant, then continued HTV for another hour. In both groups, BAL analyses indicated that LNP-XeC instillation blocked the HTV-induced BAL markers of lung inflammation and injury (Figs. 7a-d, 8a-d). LNPs that encapsulated PBS were not protective. Taking our findings together, we conclude that given prior to, or after initiation of HTV ventilation, LNP-XeC therapy protected against HTV-induced lung injury.

## DISCUSSION

Our goal was to test the commonly held view that stretch-induced AM activation underlies the lung’s mechanosensitive immune response (27). Contrary to this view however, our findings indicated that alveolar expansion caused no dimensional changes in sessile AMs. To determine whether non-uniformity of alveolar expansion was a factor in these responses (9), we interrogated the pattern of segmental shape changes in alveoli that contained AMs. These studies affirmed that while hyperinflation distended several segments of the alveolar wall, segments that contained sessile AMs did not distend, hence the AMs did not stretch. Similar responses were evident in human and pig lungs that also contained alveolus adherent AMs in non-distending alveolar segments. Together, these findings revealed the new understanding that sessile AMs located at non-distending alveolar niches were strain protected in that they did not stretch during alveolar overexpansion.

These interpretations must be made with caution since our RCM findings were obtained in subpleural alveoli that may expand differently from alveoli in deeper lung regions. However, we note that although regional variation of lung expansion depends on variation in the transpulmonary pressure, alveoli themselves expand similarly at subpleural and deeper lung levels as reported in classical studies (28). Given this similarity, we anticipate that strain heterogeneities and AM location in strain protected alveolar segments occurs in all alveoli.

However, differences between mouse, human and pig lungs may not have been adequately captured in our experiments, and require further study.

The increase of AM Ca^2+^ emerged as the major hyperinflation-induced immunogenic mechanism. We replicated this response in isolated and in HTV ventilated lungs, indicating that AM Ca^2+^ increases occurred for both a single as well as for multiple hyperinflations. The single hyperinflation model revealed that the Ca^2+^ response was prolonged in AMs despite the brief hyperinflation, a time course we reported previously in alveolar epithelium (19). By contrast in cultured alveolar type 2 cells, stretch-induced Ca^2+^ transients are short (29, 30), possibly because the cells lack GJC, hence Ca^2+^ communication with adjoining alveolar type 1 cells. In isolated lungs, the AM Ca^2+^ increases induced TNFα secretion and epithelial loss of TNFR1, signifying initiation of TNFα-induced immune signaling (22). These findings were replicated in HTV ventilated lungs, affirming that findings in the single hyperinflation and HTV models were similar.

The puzzle of how the Ca^2+^ increases occurred despite lack of AM stretch was resolved through FRAP assays that revealed a subset of AMs that formed GJC with the adjacent alveolar epithelium. Inhibition of the GJC by *Cx43* deletion in AMs blocked the hyperinflation-induced responses in AMs. Further, AM-targeted LNP-XeC delivery blocked both Ca^2+^ increase and TNFα secretion in AMs. Taking these findings together, we conclude that hyperinflation activated Ca^2+^ communication between the alveolar epithelium and AMs across Cx43-dependent gap junctional channels, resulting in TNFα secretion. Hence, the presence of Cx43 in AMs, but not AM stretch, critically determined the lung’s mechanosensitive immune response to hyperinflation.

The importance of this Cx43 based mechanism as a determinant of the lung’s global mechano-immune response was revealed by mechanically ventilating mice at HTV. These mice developed marked increases in the neutrophil count as well as in the levels of proinflammatory cytokines and plasma proteins in the BAL, affirming reported findings (31). However, AM-specific *Cx43* deletion blocked all these responses. Thus, our findings in mechanically ventilated lungs were similar to those in isolated lungs, in that both models GJC between the epithelium and AMs was revealed as the critical injury-initiating mechanism. We speculate that Ca^2+^-induced TNFα secretion from AMs followed by ligation of TNFR1 on the alveolar epithelium induced a sequence of events that led to secretion of multiple cytokines, causing lung injury as reflected by increased barrier permeability to plasma proteins.

The burgeoning interest in AM-specific drug delivery addresses the critical role played by AMs in initiating and sustaining lung injury (32). Surfactant given alone is ineffective in improving mortality in mechanically ventilated patients with ARDS (33). However, we show here that surfactant might be an effective vehicle for AM therapy as it mediated AM-specific delivery of LNPs. Notably, surfactant-associated LNPs were not taken up in the alveolar epithelium, or in systemic organs, ruling out the possibility of potential non-specific effects. By contrast, given without surfactant LNPs were taken up non-specifically as we show here, and others report (34). LNPs have been directed at specific AM receptors (35). However, such receptor targeting might access only those AMs that express the receptor. Our strategy takes advantage of the fact that all AMs internalize surfactant (36). Hence, by present strategy surfactant-associated LNPs are likely to target all AMs. Accordingly, the instillation of surfactant-associated LNP-XeC blocked the hyperinflation-induced Ca^2+^ increase in AMs, as well as the lung injury response to HTV. To our knowledge, our findings indicate airway instillation of surfactant-associated LNPs provides a therapeutic platform for AM-specific targeting.

An important set of findings relates to the comparison of responses in mouse, human and pig alveoli. Although such comparisons are likely to advance translational understanding, a caveat is that human lungs are transplant rejects and are therefore not pristine, potentially complicating comparisons against findings in rodent models. We addressed these difficulties through studies in pig alveoli, since pig lungs are pristine, of similar size as human lungs, and they provide a translational model for studies of mechanical ventilation (11, 12). The failure of hyperinflation to cause shape changes in AMs in mouse and pig alveoli, as also in human alveoli, suggests that our interpretation that AMs are strain protected is translationally valid.

In conclusion, our findings extend micromechanical understanding of the complex strain landscape of the distended alveolar wall. What is now clear in a translationally relevant manner, is that the non distending segment of the alveolus forms the key protective niche for sessile AMs, obviating the need to posit AM stretch as the critical mechanism underlying immune activation in mechanical ventilation. Nevertheless, the AMs formed the hub for the proinflammatory signaling responsible for tissue injury during ventilatory lung overexpansion, since proinflammatory Ca^2+^ signals passed from the epithelium to the AMs across gap junctions. The consequences were critical, as inhibition of the signal in an AM-targeted manner suppressed lung inflammation and injury. The extent to which similar therapy might apply to other AM-initiated inflammatory lung diseases requires consideration.

## MATERIALS AND METHODS

### Materials

A list of antibodies used in this study is available in the supplementary material (Table. s1). We purchased mAb MCA2350 against the TNFR1 extracellular epitope (40 μg/ml) from AbD Serotec (Oxford, UK). Anti-TNFR1 antibody was fluorescently labeled with Alexa Fluor 633 using our standard protocol (37). Fluo-4 AM (5 µM), Lysotracker Red (LTR, 100 nM), calcein AM (5 µM), FM1-43 (5 µM) and rhodamine-B labelled dextran 70 kDa (80 μg), were purchased from Thermo-Fisher Scientific (Waltham, MA). Xestospongin C (XeC, 20 μM) was purchased from Sigma-Aldrich (St. Louis, MO). We purchased human recombinant TNFα (10 ng/ml) from BD Biosciences (San Jose, CA). Surfactant (CUROSURF®, poractant alfa, Chiesi Farmaceutici (Parma, Italy)) was used at a final concentration of 5 mg/ml. FM1-43 dye (5 µM) and surfactant (5 mg/ml) were incubated together away from light for 10 min at room temperature to generate fluorescent (FM1-43 labelled) surfactant. Vehicle for fluorophores, antibodies and other agents was HEPES buffer (see below for composition).

### Mouse care

All mouse procedures were reviewed and approved by the institutional animal care and use committee (IACUC) at Columbia University Irving Medical Center. All mice were cared for according to the National Institutes of Health (NIH) guidelines for the care and use of laboratory animals. Mice were socially housed under a 12 h light/dark cycle with ad libitum access to water and food. The mice were inbred hence no randomization was required.

Nevertheless, we randomly assigned age- and sex matched mice to experimental groups.

Mice were between 2–4 months and on a C57BL/6J background. CD11cCre^+/-^ mice were provided by Boris Reizis while Cx43^flox/flox^ (stock no. 008039) and C57BL/6J (stock no. 000664) mice were purchased from the Jackson Laboratory and bred in house.

### Isolated, blood perfused mouse lung preparation

Using our reported methods (16), we anesthetized (I.P., ketamine 100 mg/kg, xylazine 10 mg/kg) and heparinized mice (1000 IU/kg, intra-cardiac). Then we exsanguinated mice and excised the lungs for pump perfusion (0.5 ml/min, 37°C) through the pulmonary artery (PA) using autologous blood (1 ml of 5 ml total perfusate) with added HEPES buffer (150 mmol/l Na^+^, 5 mmol/l K^+^, 1.0 mmol/l Ca^2+^, 1 mmol/l Mg^2+^, and 20 mmol/l HEPES, pH 7.4) containing 70 kDa dextran (4% w/v) and 1% FBS. Osmolarity was 300 mosM (Fiske Micro-Osmometer, Fiske® Associates, Norwood, MA). The left atrial outflow was recirculated through the PA. Lungs were inflated with room air through a tracheal cannula. We held the pulmonary artery, left atrial and airway pressures at 10, 3 and 5 cmH_2_O respectively during microscopy (sFig. 8).

**Figure 8.**
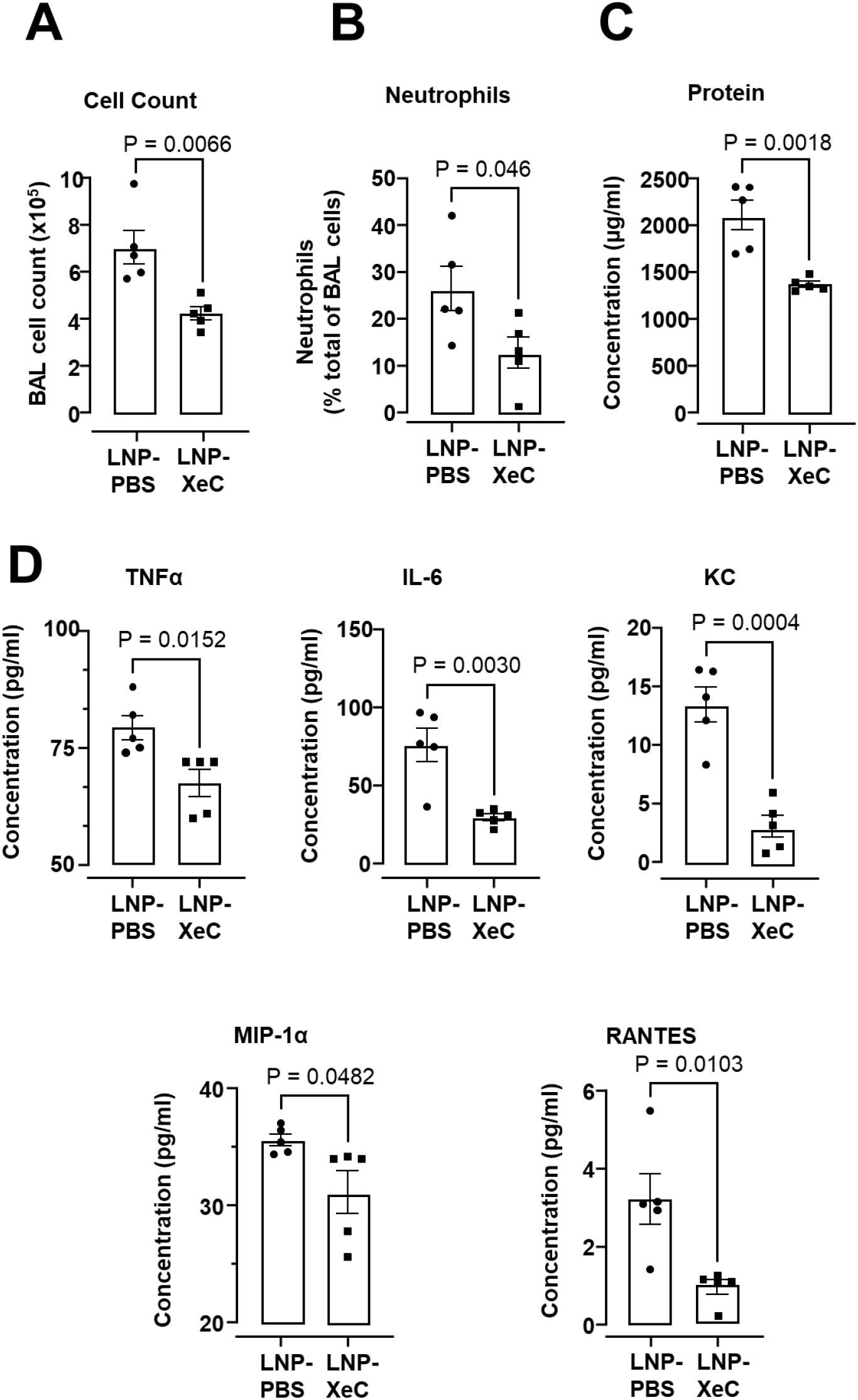
LNP-XeC given therapeutically attenuates high tidal volume ventilation- induced lung injury **a-d** Bars show quantifications in the bronchoalveolar lavage (BAL) for total cells (a) neutrophil content (b), protein concentration (c), cytokine and chemokine levels (d) in mice mechanically ventilated for 1 hour at 18 ml/kg then instilled with lipid nanoparticles (LNPs) encapsulating XeC or PBS in a surfactant solution then ventilation continued for 1 hour. LNP-*PBS*, mice instilled with LNPs encapsulating PBS, *LNP-XeC*, mice instilled with LNPs encapsulating XeC. LNPs were given in a surfactant containing solution. Mean±sem, n = 5 lungs per group. Data analyzed by unpaired t-test.

### Alveolar micropuncture and microinfusion

Using glass micropipettes (tip diameter 3-5 μm), we microinfused single alveoli with fluorescent dyes and antibodies (sFig. 8). Infusates were delivered to ∼7 alveoli around the micropuncture site (16). For each bolus, we microinfused alveoli for ∼3 s after which the liquid drained from the alveolar lumen in seconds, re-establishing air-filled alveoli. This rapid clearance indicates that the micropuncture did not rupture the alveolar wall, and that the micropunctured membrane rapidly resealed as reported for other cells (38). Nevertheless, we selected non-micropunctured alveoli for imaging. To determine whether the fluorescence detected in the epithelium was intracellular or extracellular, in some experiments we microinfused alveoli with the fluorescence quencher, trypan blue (0.01% w/v, Thermofisher, Waltham, MA). Persistent fluorescence after trypan blue wash out indicated intracellular labelling (37). In all experiments in which multiple dyes were infused, we confirmed absence of bleed-through between fluorescence emission channels.

### Lipid nanoparticle preparation

Following the manufacturers protocol, we extruded unilamellar lipid nanoparticles (LNP, 20 μg/μl; 100 nm pore size; DOTAP, Avanti Lipids, Birmingham, AL) in sterile Opti-MEM (Invitrogen, Carlsbad, CA). We used freshly extruded LNPs for experiments. Using our established protocol (37), LNPs were complexed with plasmid DNA, rhodamine-labelled dextran 70kDa, XeC or PBS depending on downstream experiment.

### Plasmid preparation and *In vivo* mouse sessile AM lung transfection

GFP-TNFα plasmid (39) (plasmid #28089) was purchased from Addgene (Watertown, MA). We transformed DH5α *E*. *coli* (New England Biolabs) with plasmid DNA via heat shock, then amplified and purified the plasmids using an EndoFree Plasmid Maxi Kit (Qiagen). Using our established methods (16), we complexed plasmid DNA (80 μg total) with lipid nanoparticles (LNP, 480 μg total) in sterile Opti-MeM. We administered a plasmid DNA–LNP mix (60 μl) in a surfactant containing suspension (5 mg/ml) by intranasal instillation (I.N.) of anesthetized mice. Imaging experiments were carried out 48 hours after transfection.

### Live imaging of mouse lungs

We imaged intact alveoli in a 1.7-μm-thick optical section (512 x 512 pixels) at a focal plane ∼20 μm deep to the pleural surface of live lungs with laser scanning microscopy (TCS SP8, Leica Microsystems, Wetzlar, Germany) using a 10x air objective (numerical aperture 0.3, Leica Microsystems, Wetzlar, Germany) or 25x water immersion objective (numerical aperture 0.95, Leica Microsystems, Wetzlar, Germany) (sFig. 8). For Z stacks, we imaged 1.7-μm-thick optical sections (512 x 512 pixels) at vertical intervals of 1.7 μm from the pleural surface to a depth of 40 μm. To immerse the 25x objective in water, we placed a water drop on a coverslip that was held in a metal O-ring, as described previously (16). We used our reported methods to detect fluorescence of microinfused dyes and antibodies in live alveoli (8). Cytosolic Ca^2+^ was detected using the cytosolic Ca^2+^ indicator, fluo-4 AM, an acetoxymethyl (AM) ester that fluoresces upon intracellular cleavage of the AM moiety as previously reported (8).

Mitochondrial Ca^2+^ was detected using the mitochondrial Ca^2+^ indicator, rhod-2-AM as previously reported (16). We recorded Ca^2+^ responses in alveolar epithelium and macrophages at one image/10 s as previously reported (8). All images were recorded as single images and processed using Image J (v15.3, National Institutes of Health). We applied brightness and contrast adjustments to individual pseudocolor channels of entire images and equally to all experimental groups. No further downstream processing or averaging was applied.

### Mouse lung hyperinflation

To induce lung hyperinflation, we increased airway pressure from 5 to 15 cmH_2_O for 15 seconds. After 15 seconds we reduced airway pressure to 5 cmH_2_O and identified the pre- hyperinflation optical section using morphological landmarks as previously described (9). To confirm alveolar stretch due to hyperinflation in mouse lungs, we recorded the time-dependent secretion of surfactant by alveolar epithelial type II cells as loss of lysotracker red (LTR) fluorescence post hyperinflation as shown (sFig. 9) and as previously described (19).

**Figure 9.**
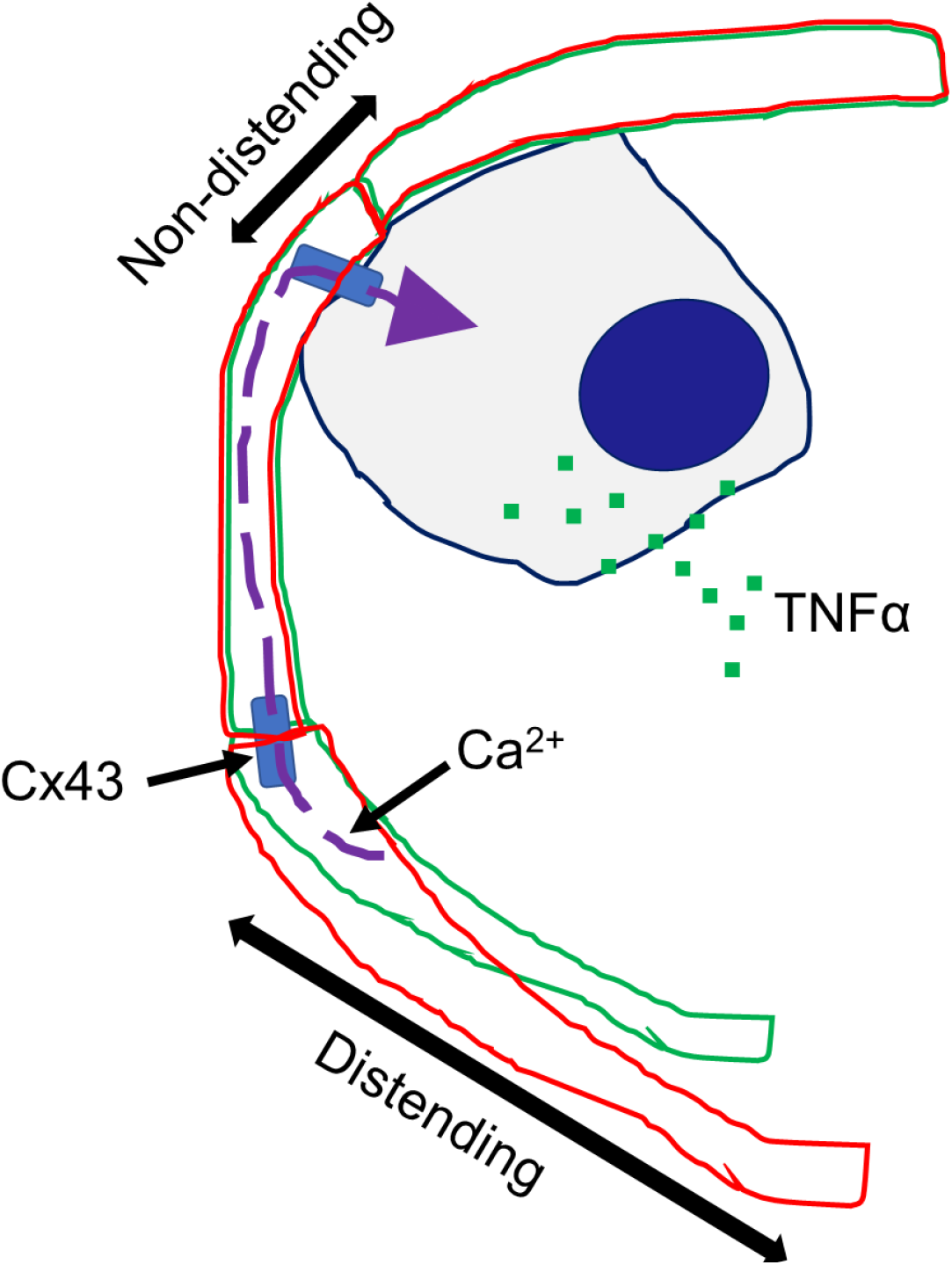
Ca^2+^ communication between distending and non-distending alveolar segments. Alveolar overexpansion due to lung hyperinflation causes distention of AM-free alveolar segments. Ca^2+^ increase in the distended segment diffuses intracellularly through Cx43 gap junctions to AMs in the non-distended segment. The Ca^2+^ increase in AMs initiates TNFα secretion. *Cx43*, connexin-43.

### Live imaging of pig lungs

Confocal microscopy on live pig lungs was carried out in lungs obtained from 2 male and 1 female (3-month-old) Yorkshire strain pig donors. Pigs were anaesthetized by intramuscular injection of ketamine (100 mg/kg) and xylazine (10 mg/kg). The lungs were removed immediately after the pig was euthanized by potassium chloride overdose by intracardiac injection. The lingular lobe, which provides a flat surface convenient for live microscopy, was positioned below the objective of a two-photon microscope (TCS SP8, Leica). The lobe’s pulmonary artery pressure was held at 10 cmH_2_O, while the lobe was inflated at alveolar pressure of 5 cmH2O through a bronchial cannula. We injected fluorescent dyes and antibodies through a 31-gauge needle into alveoli. To induce alveolar expansion, lungs were inflated by increasing airway pressure from 5 to 15 cmH_2_O. For a single hyperinflation protocol, airway pressure was increased from 5 to 15 cmH_2_O for 15 seconds.

### Live imaging of human lungs

Confocal microscopy on live human lungs was carried out in lungs obtained from 3 male and 1 female (ages 34 – 82) de-identified donors. The lungs were imaged after ∼20 h of cold ischemia. Human lungs were warmed with perfusion of 37 °C buffer prior to and during imaging. The temperature of the lung’s venous outflow was directly measured and set at 37 °C which represented the lung’s core temperature. The lingular lobe, which provides a flat surface convenient for live microscopy, was positioned below the objective of a two-photon microscope (TCS SP8, Leica). The lobe’s pulmonary artery pressure was held at 10 cmH_2_O, while the lobe was inflated at alveolar pressure of 5 cmH_2_O through a bronchial cannula. We injected fluorescent dyes and antibodies through a 31-gauge needle into alveoli. To induce alveolar expansion, lungs were inflated by increasing airway pressure from 5 to 15 cmH_2_O. For a single hyperinflation protocol, airway pressure was increased from 5 to 15 cmH_2_O for 15 seconds. Human lungs were rejected for experiment if: (1) there was presence of exudates and cells denoting alveolar disease, and (2) there was a failure to detect alveolar expansion following a hyperinflation pressure challenge.

### Fluorescence recovery after photobleaching

In RCM studies of live mouse lungs, we quantified fluorescence recovery after photobleaching (FRAP), which assays gap junctional communication (GJC) as per our previous protocol (8, 40). We first identified sessile AMs (SiglecF^+^ cells) in which the Ca^2+^ or TNFα secretory response was present or absent then we determined GJC by FRAP. We photobleached cytosolic fluorescence within AMs and determined fluorescence recovery indicating intercellular dye transfer and GJC with alveolar epithelium. Lack of fluorescence recovery indicates lack of GJC.

### Intranasal and intratracheal instillation

For intranasal instillation, mice were anaesthetized (I.P., ketamine 50 mg/kg, xylazine 5 mg/kg) then instilled with 2 ml/kg of instillate and allowed to recover for 2 hours prior to the experiment. For intratracheal instillation during mechanical ventilation protocol, ventilation was paused by removing the tracheal cannula from the ventilator then anaesthetized mice were instilled with 2 ml/kg of instillate through the cannula. The cannula was reconnected to the ventilator after instillation. For LNP-XeC instillation, mice were I.N. instilled with 20 µM XeC complexed in LNPs (20 μg/μl) suspended in a surfactant-PBS (Ca^2+^ and Mg^2+^ free) suspension (5 mg/ml). 60 μl of LNP-XeC suspension was instilled.

### Mechanical ventilation

A previously described mechanical ventilation (MV) protocol with modifications was used (25). Briefly, mice were anesthetized with isoflurane (4%) and ketamine (50 mg/kg) and xylazine (5 mg/kg) administered I.P. A tracheal cannula (PE-90 tubing, BD, Franklin Lakes, NJ) was inserted and secured into place by suture, and MV was started with a tidal volume of 6 ml/kg, a positive end-expiratory pressure (PEEP) of 5 cmH_2_O and a respiratory rate of 120 breaths/min (VentElite, Harvard Apparatus, Holliston, MA). Following a 10-min stable baseline period, in some experiments tidal volume was incrementally increased to 18 ml/kg over 5 minutes for high tidal volume ventilation (HTV) or continued at 6 ml/kg for low tidal volume ventilation (LTV). Depth of anesthesia was assessed by reaction to paw pinch throughout the protocol.

Anesthesia was maintained with ketamine (20 mg/kg) and xylazine (2 mg/kg) both administered by I.P injection. MV was continued for 2 hours. Saline (2 ml/kg) was administered in 30 min intervals by I.P injection throughout the protocol. Blood oxygen saturation, airway pressures, blood pressure, and heart rate were monitored continuously using a computer-integrated data collection system (MouseOx Plus, Starr Life Sciences, Oakmont, PA). Temperature was monitored using a rectal probe and temperature was maintained at 37 °C throughout the protocol using a heating blanket (Homeothermic Blanket Control Unit, Harvard Apparatus, Holliston, MA). After 2 hours of MV, BAL was performed for analysis of BAL fluid and ALI was evaluated.

### Mechanical ventilation of isolated, perfused mouse lungs

We setup isolated, perfused lungs as described above. We microinfused relevant fluorescent agents as described above and carried out real-time fluorescence imaging for baseline recordings. For mechanical ventilation of isolated, perfused lungs the tracheal cannula connected to a continuous positive airway pressure (CPAP) machine was reconnected to a mechanical ventilator (VentElite, Harvard Apparatus, Holliston, MA). MV was initiated with a tidal volume of 6 ml/kg, a positive end-expiratory pressure (PEEP) of 5 cmH_2_O and a respiratory rate of 120 breaths/min (VentElite, Harvard Apparatus, Holliston, MA). Following a 5-min stable baseline period, tidal volume was incrementally increased to 18 ml/kg over 5 minutes for high tidal volume ventilation (HTV) for 2 hours. After 2 hours of HTV, the tracheal cannula was reconnected to the CPAP machine and imaging of lungs was immediately continued for post HTV recordings.

### Analysis of BAL fluid

The lungs of mice were lavaged repeatedly (5x) with the same 1 ml ice-cold PBS (Ca^2+^ and Mg^2+^ free) through a tracheal cannula. Collected BAL fluid was centrifuged for 4 minutes at 400 g at 4°C. The supernatant was analyzed for protein and cytokine concentrations. We measured TNFα, IL-6, KC (CXCL1), MIP-1α (CCL3) and RANTES (CCL5) concentrations in BAL fluid to determine pro-inflammatory cytokine secretion. Sample testing was carried out by Quansys Biosciences using a multiplex chemiluminescence assay (Q-plex) for the detection of mouse cytokines.

### Evaluation of ALI

We evaluated ALI by assessing BAL protein content by BCA assay (Thermo Fisher, Waltham, MA) to determine protein leak from vascular compartment into alveolar airspace. We also assessed ALI by characterizing cell populations in BAL by flow cytometry (see below).

### Flow cytometry analysis of BAL cells

Briefly, BAL cell pellet was resuspended in PBS (Ca^2+^ and Mg^2+^ free) supplemented with 1% FBS, 100 mM HEPES. and used for flow cytometry studies. Relevant antibodies were added to the cell suspension and incubated away from direct light for 1 hour at room temperature. The cells were centrifuged (4 min, 4 °C, 400 g) and resuspended in buffer. This procedure was repeated two more times, and the cells were finally resuspended in 300 ul PBS (Ca^2+^ and Mg^2+^ free) without supplements. We analyzed cells by flow cytometry (3L Cytek Aurora, Becton Dickinson, NJ) according to manufacturer’s protocols using standard software (FCS Express 7 Flow, De Novo Software). Neutrophils were identified as CD45-Pacific Blue, Ly6G-PE-Cy7, and CD11b-PeCy5 positive cells following a standard gating protocol (sFig. 10) (41).

### Flow cytometry analysis of LNP uptake by lung cells

Mice were I.N. instilled with 80 μg rhodamine-B labelled dextran 70 kDa (Thermo Fisher, Waltham, MA) complexed in LNPs (480 μg total) suspended in a surfactant containing suspension (5 mg/ml). 60 μl of LNP-surfactant suspension was instilled. 2 hours later, we isolated lung cells. To isolate cells, we buffer (PBS, Ca^2+^ and Mg^2+^ free) perfused lungs through vascular cannulas to clear blood; we then minced and passed the tissue through 40 μm cell strainers (BD Biosciences, Franklin Lakes, NJ) to obtain a single cell suspension. For flow cytometry, we surface stained the cells by incubating the suspension with relevant antibodies away from direct light for 1 hour at room temperature. We analyzed cells by flow cytometry (5L Cytek Aurora, Becton Dickinson, Franklin Lakes, NJ) according to manufacturer’s protocols using standard software (FCS Express 7 Flow, De Novo Software). Alveolar macrophages were identified as SiglecF-PE, and CD11c-APC positive cells following a standard gating protocol (sFig. 11) (41). Alveolar epithelial cells were identified as EpCam- PerCP-Cy5.5 positive cells.

### Flow cytometry analysis of LNP uptake by heart, spleen and liver cells

Mice were I.N. instilled with 80 μg rhodamine-B labelled dextran 70 kDa (Thermo Fisher, Waltham, MA) complexed in LNPs (480 μg total) suspended in a surfactant containing suspension (5 mg/ml). 60 μl of LNP-surfactant suspension was instilled. 2 hours later, we isolated heart, spleen and liver cells. To isolate cells, we buffer (PBS, Ca^2+^ and Mg^2+^ free) perfused systemic circulation through vascular cannulas to clear blood; we then minced and passed the tissues through 40 μm cell strainers (BD Biosciences, Franklin Lakes, NJ) to obtain single cell suspensions. For flow cytometry, we surface stained the cells from each organ by incubating the suspension with relevant antibodies away from direct light for 1 hour at room temperature. We analyzed cells by flow cytometry (5L Cytek Aurora, Becton Dickinson, Franklin Lakes, NJ) according to manufacturer’s protocols using standard software (FCS Express 7 Flow, De Novo Software). LNPs were identified as rhodamine B positive events following a standard gating protocol (sFig. 3).

### Human lung study approval

As confirmed by the Columbia University IRB, per the NIH policy, since all samples in the study were acquired from deceased individuals, the study was not considered human subject research. This policy is based on United States Department of Health and Human Services human subject regulations under 45 CFR 46 wherein a human subject is defined as “a living individual about whom an investigator (whether professional or student) conducting research obtains (1) data through intervention or interaction with the individual or (2) identifiable private information”.

### Participants

Confocal microscopy was done with intact human lungs obtained from brain dead organ donors at the time of tissue acquisition for transplantation and when not used for clinical transplant, as described (10) through collaboration and protocol with LiveOnNY, the organ procurement organization for the New York area. Demographic data are detailed in Supplementary Table 2.

### Patient consent

In every case, patient surrogates (individuals with power of attorney/next of kin) provided written consent for the use of deceased donor organs in research. Research, including consent for research, was conducted in alignment with the Declaration of Helsinki.

### Privacy protection

None of the investigators had access to identifiable private information, and all samples were assigned unique, non-identifying IDs on receipt. Their correspondence to the United Network for Organ Sharing (UNOS) IDs assigned by the organ procurement organizations (OPOs) and LiveOnNY was not communicated to the OPOs by the investigators, and they are maintained by the investigators under a secure, password-protected network.

### Statistics

All groups comprised a minimum of 3 mice each. Group numbers were designed to enable detection of statistically significant differences with a power of at least 85%. For applicable imaging experiments, we carried out a paired protocol, where baseline and test conditions were obtained in the same alveolus or cell, and at least 3 determinations were obtained per lung. These determinations were averaged to obtain a mean for each condition in each lung. The means for each lung were pooled for the group to obtain mean±sem, where n represents the number of lungs unless otherwise stated. Data are shown as mean±sem. We analyzed paired data by the paired t-test and non-paired data by unpaired t-test. Significance was accepted at P < 0.05.

## List of Supplementary Materials

Supplementary Figures 1 to 11.

Supplementary Tables 1 to 2.

## Acknowledgments

Human lungs were obtained from an organ procurement organization (LiveOnNY) with support from Dr. Donna Farber, Department of Surgery, Columbia University. The Flow Cytometry Core was supported by NIH award S10OD020056.

## Author contributions

Conceptualization: LM, JB Methodology: LM, MNI, GAG, BK Investigation: LM, MNI Visualization: LM, JB Funding acquisition: LM, JB Project administration: LM, JB Supervision: JB Writing – original draft: LM Writing – review & editing: LM, MNI, SB, JB The order of co–first authors was determined based on LM’s greater contribution to conceptualization, methodology, investigation, visualization, and writing.

## Data and materials availability

All data are available in the main text or the supplementary materials.

