## Supplemental files for "Macrophage-specific lipid nanoparticle therapy blocks the lung’s mechanosensitive immunity due to macrophage-epithelial interactions"

**This file includes:**

Supplementary Figures 1 to 11
Supplementary Tables 1 to 2

#### SUPPLEMENTARY FIGURE 1

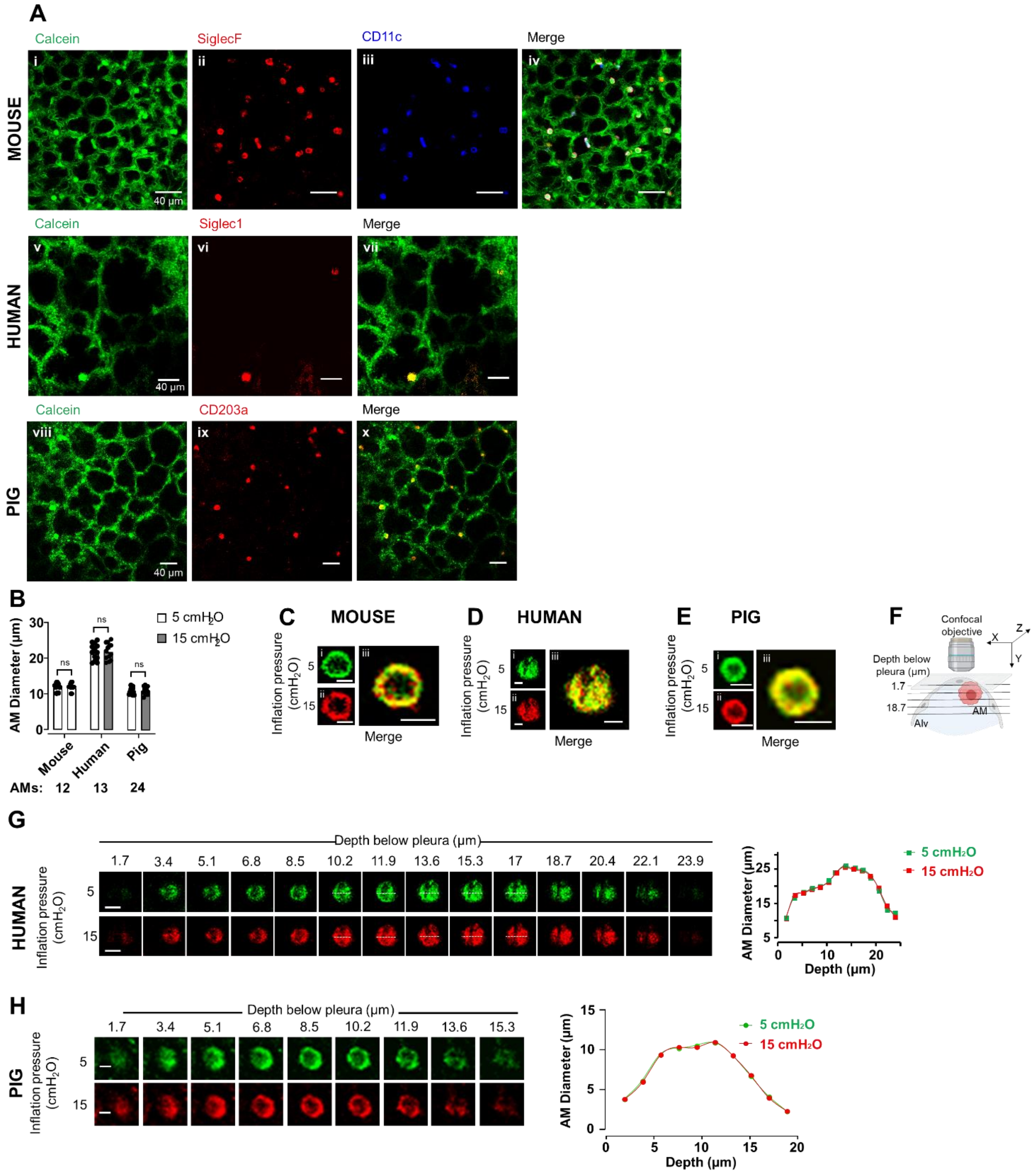


**Supplementary Figure 1. Sessile AM real time confocal microscopy**

**a** Confocal images (*i-iv*) show calcein loaded alveoli from mouse lung. AMs are identified by SiglecF and CD11c fluorescence. Confocal images (*v-vii*) show calcein loaded alveoli from human lung. AMs are identified by Siglec1. Confocal images (*viii-x*) show calcein loaded alveoli from pig lung. AMs are identified by CD203a.

**b** Bars show paired diameter quantifications for single macrophages (dots) at the indicated inflation pressures in mouse, human and pig lungs. Mean±sem. n = 4-6 lungs per group. Scale bars = 10 µm.

**c-e** Confocal images show a single AM from mouse (b), human (c) and pig (d) lungs at high magnification (*i-iii*). The AM immunofluorescence for SiglecF (mouse) Siglec1 (human) and CD203a (pig) are shown in green *(i)* and red *(ii)* pseudocolors for the indicated inflation pressures. AM distension was assessed by pseudocolor superimposition (*iii*). Bars show paired diameter quantifications for single macrophages (dots) at the indicated inflation pressures. Mean±sem. n = 4-6 lungs. Scale bars = 10 µm.

**f** Sketch shows the imaging protocol for g,h and Fig. 1f. Sketch was made using Biorender.

**g** Confocal z stack images show the human AM perimeter (Siglec1 staining) at 5 (green) and 15 (red) $cmH_2O$ inflation pressure at different depths. The graph plots AM diameter (dashed line) at different depths below the pleura. Replicated in 13 AMs from 3 lungs. Scale bar = 20 µm.

**h** Confocal z stack images show the pig AM perimeter (CD203a staining) at 5 (green) and 15 (red) $cmH_2O$ inflation pressure at different depths. The graph plots AM diameter (dashed line) at different depths below the pleura. Replicated in 10 AMs from 4 lungs. Scale bar = 5 µm.

#### SUPPLEMENTARY FIGURE 2

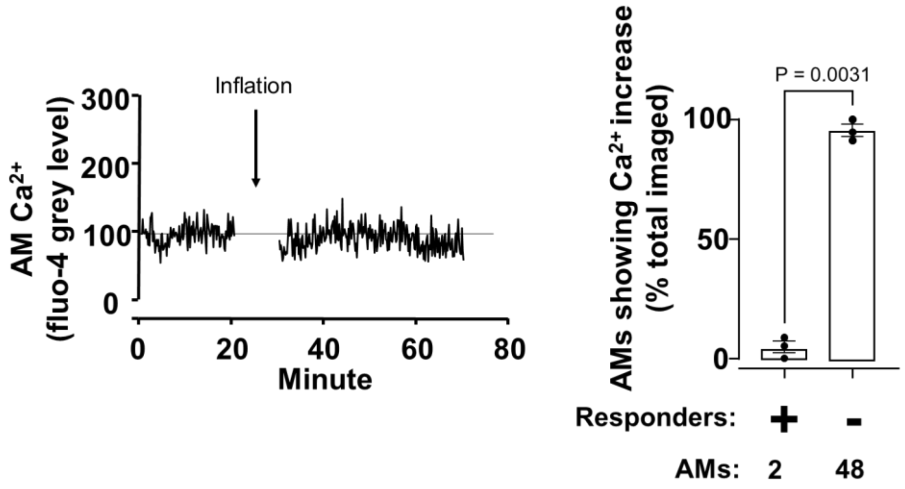


**Supplementary Figure 2. Lack of $Ca^{2+}$ increases in AMs in response to inflation from 5 to 10 $cmH_2O$**

Tracing shows alveolar macrophage (*AM*) cytosolic calcium ($Ca^{2+}$) responses before and after inflation from 5 to 10 $cmH_2O$. Black line indicates mean cytosolic calcium. Bars show lack of AM responders to 5 $cmH_2O$ inflation. Mean±sem, n = 3 lungs per group.

**SUPPLEMENTARY FIGURE 3**

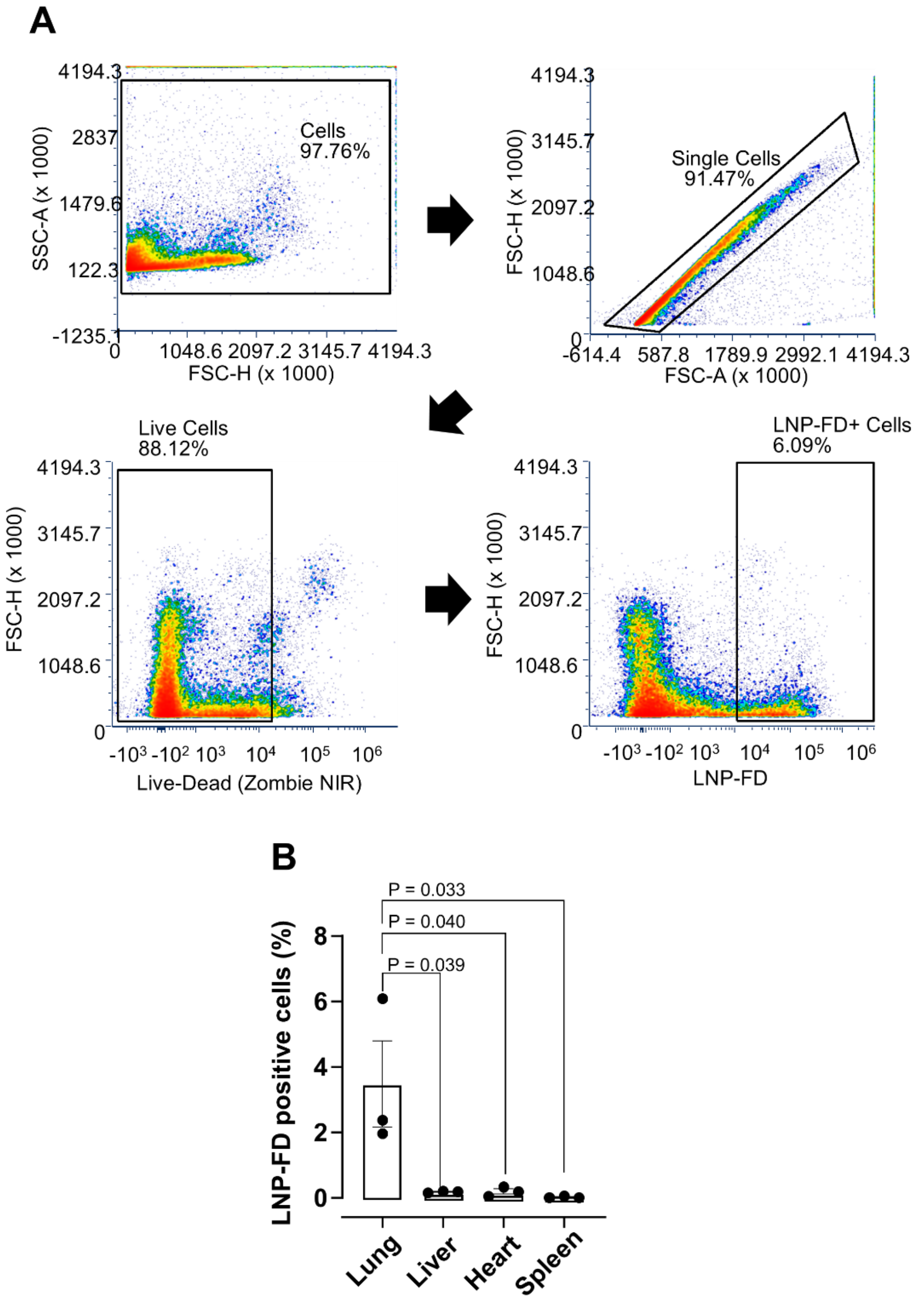


**a** Gating strategy used to identify lipid nanoparticle-encapsulating fluorescent dextran 70 kDa (*LNP-FD)* positive cells from mouse lung instilled with LNP-FD 2 hours prior. Same gating strategy was used to identify LNP-FD cells from liver, heart and spleen tissue. Cells were isolated from organs and after the exclusion of dead cells, debris and doublets, LNP-FD cells were identified by LNP-FD staining.

**b** Bars show percentage of LNP-FD positive cells in lung, liver, heart and spleen tissue from mice instilled with LNP-FD two hours prior. Mean±sem, n = 3 per group.

#### SUPPLEMENTARY FIGURE 4

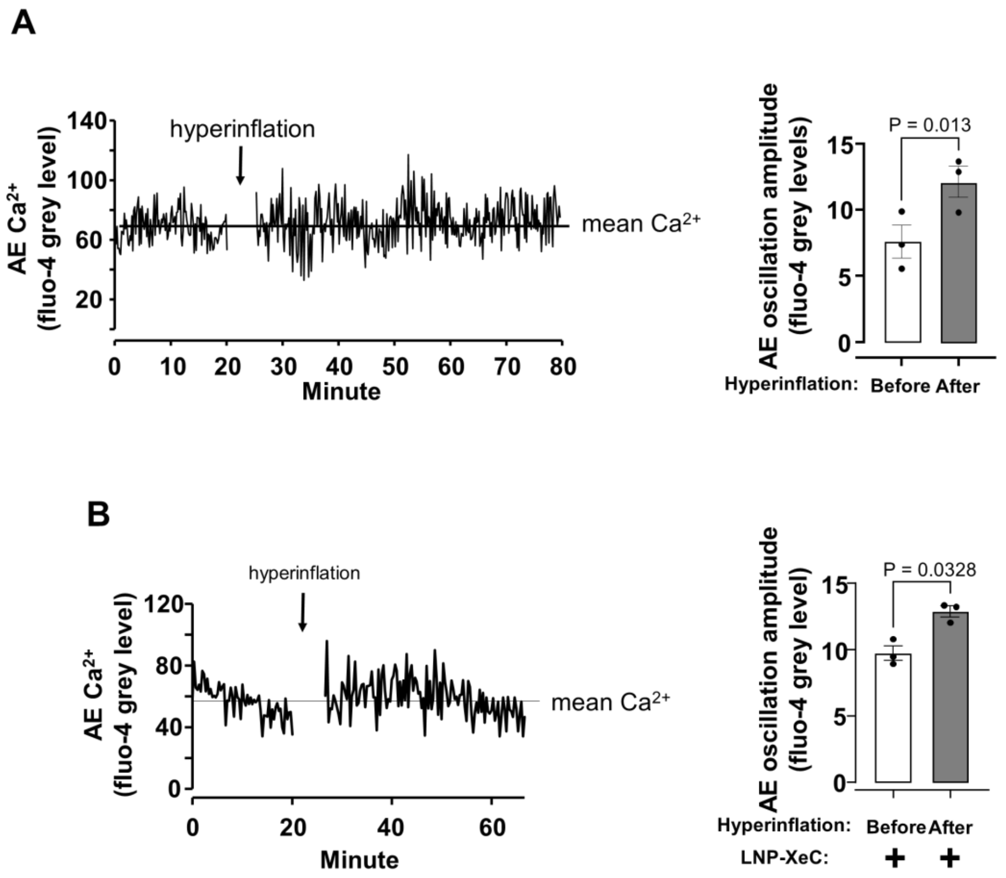


**Supplementary Figure 4. Hyperinflation-induced increases in alveolar epithelial $Ca^{2+}$ oscillations in lungs treated with or without LNP-XeC**

**a** Tracing shows alveolar epithelial (AE) cytosolic calcium ($Ca^{2+}$) responses before and after hyperinflation. Black line indicates mean cytosolic calcium. Bars show paired quantifications of AE $Ca^{2+}$ oscillation amplitude in AE (dots) before and after hyperinflation. Mean±sem, n = 3 lungs per group.

**b** Tracing shows alveolar epithelial (AE) cytosolic calcium ($Ca^{2+}$) responses before and after hyperinflation in lungs instilled with lipid nanoparticle-encapsulating Xestospongin C (LNP-XeC) 2 hours prior to experiment. Black line indicates mean cytosolic calcium. Bars show paired quantifications of AE $Ca^{2+}$ oscillation amplitude in AE (dots) before and after hyperinflation. Mean±sem, n = 3 lungs per group.

#### SUPPLEMENTARY FIGURE 5

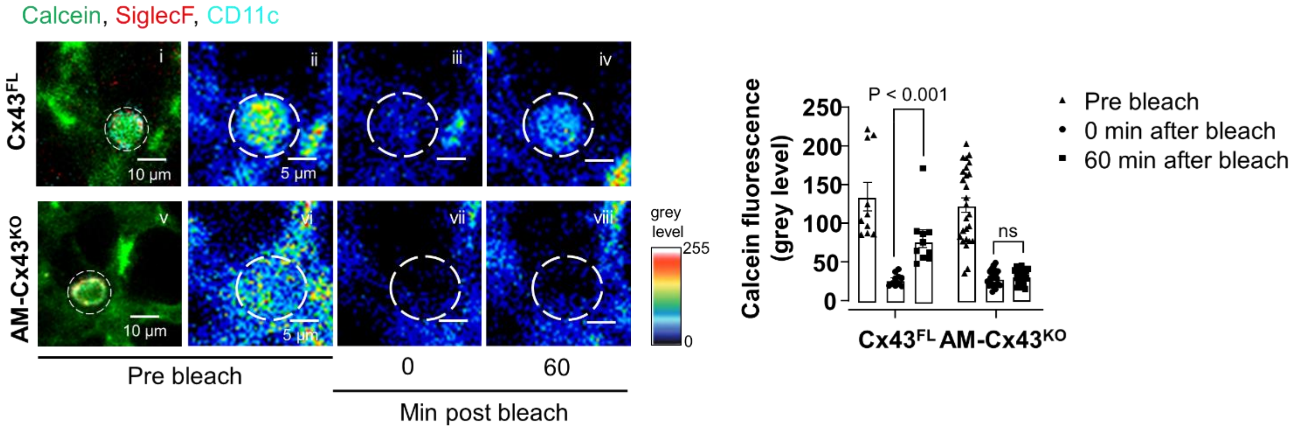


**Supplementary Figure 5. Sessile AM Cx43 deletion inhibits gap junctional communication**. Confocal images (*i, v*) show calcein green-loaded sessile AMs identified by SiglecF and CD11c labeling. Pseudocolor high power images of sessile AMs in (*i, v*) pre-bleach, 0 and 60 min after bleach (*ii*-*vii*). White dashed circles indicate photobleached regions. Bars show group data quantifications of calcein fluorescence within sessile AMs at indicated time points. Mean±sem, n = 3 lungs, 10 sessile AMs (*$Cx43^{FL}$*), n = 4 lungs, 26 sessile AMs (*$AM\text{-}Cx43^{KO}$*).

### SUPPLEMENTARY FIGURE 6

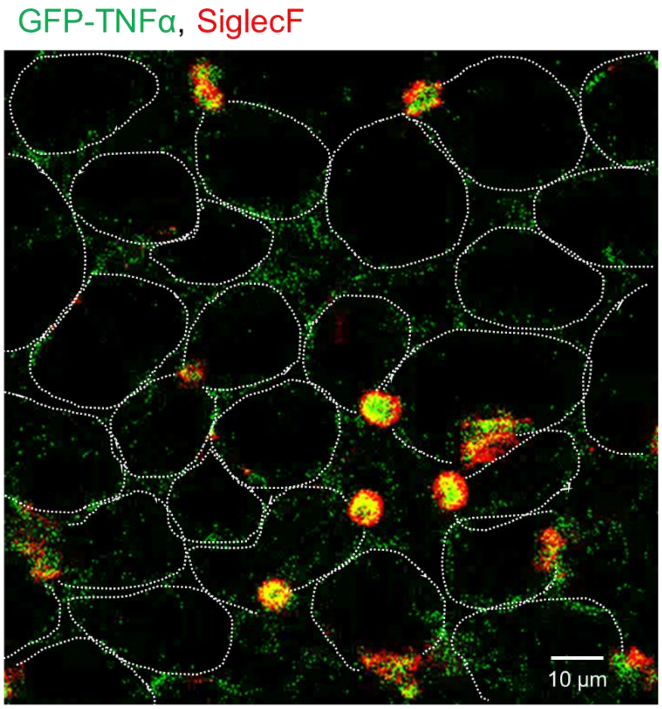


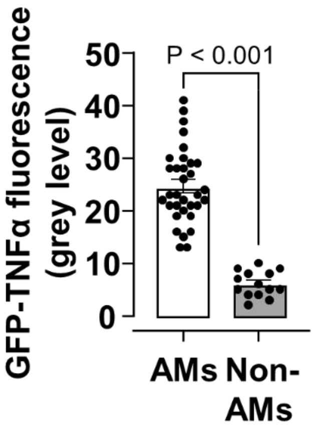


**Supplementary Figure 6. GFP-TNFα fluorescence in sessile AMs**

Confocal image shows a field of AMs which were transfected with lipid nanoparticles containing GFP-TNFα plasmid in a surfactant containing solution by intranasal instillation two hours prior. Bars shows that majority of TNFα fluorescence expressed in imaged sessile AMs compared to randomly selected non-AM (SiglecF negative) regions. White dashed lines indicate alveolar perimeters. Mean±sem, n = 3 lungs, 29 AMs. *AM*, sessile AM.

### SUPPLEMENTARY FIGURE 7

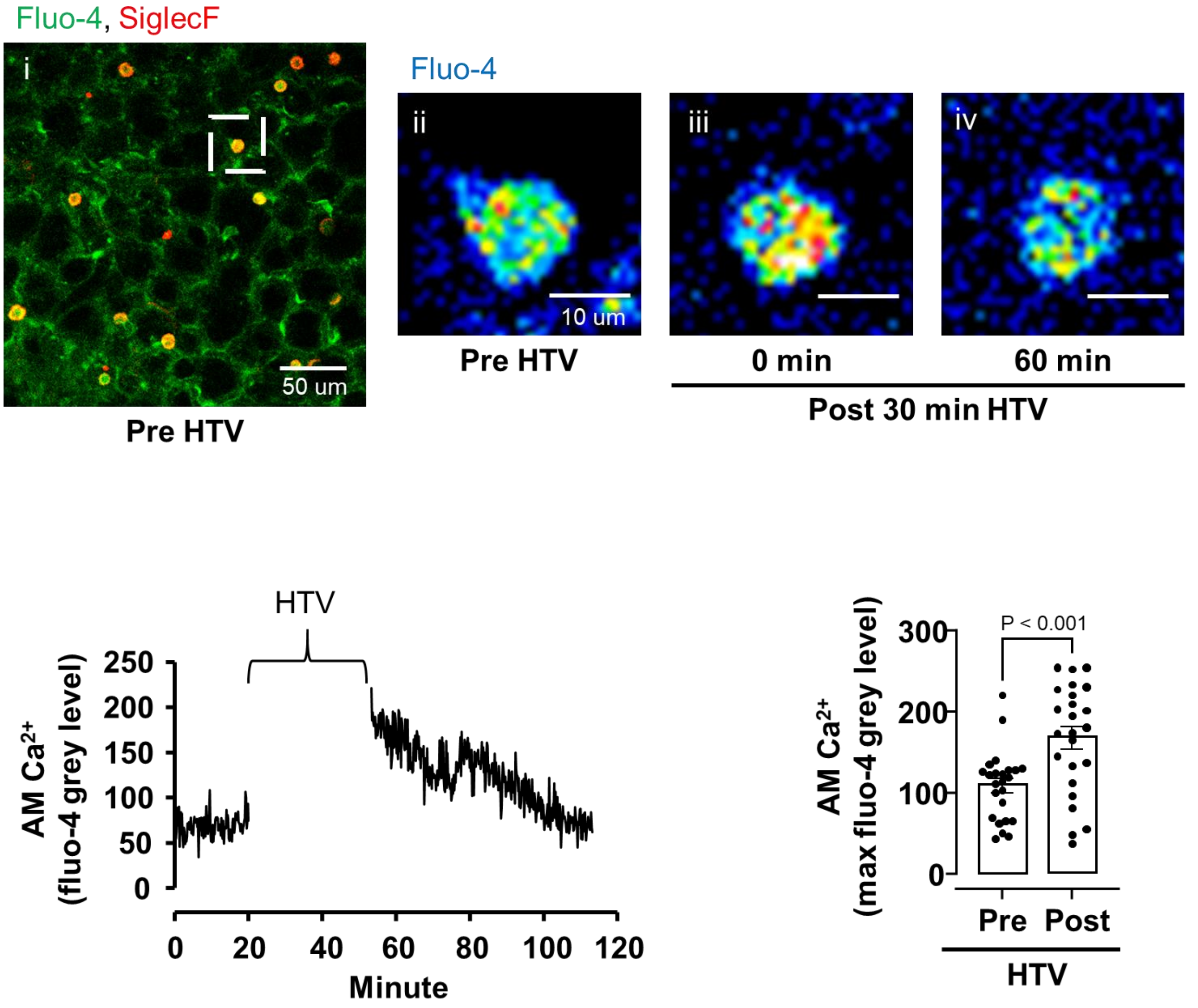


**Supplementary Figure 7. High tidal volume mechanical ventilation mobilizes $Ca^{2+}$ in sessile AMs**

Confocal image (*i*) shows $Ca^{2+}$ in sessile AMs. High magnification confocal images (*ii*-*iv*) show $Ca^{2+}$ in AM (square in *i*) before and after 30 min of high tidal volume mechanical ventilation (HTV). Tracing shows AM $Ca^{2+}$ response before and after HTV. Bars show paired comparisons of maximum $Ca^{2+}$ pre- and post-HTV in AMs. n = 24 AMs from 3 lungs.

### SUPPLEMENTARY FIGURE 8

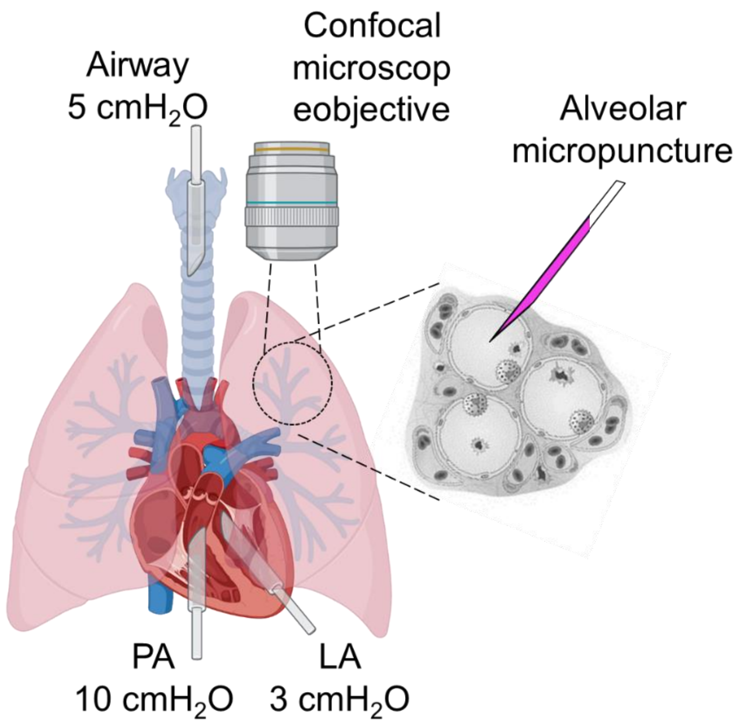


**Supplementary Figure 8 – Isolated, perfused lung preparation and alveolar micropuncture technique.** Schematic of isolated, perfused lung preparation and alveolar micropuncture technique to deliver fluorescent agents that permit visualization of the alveolar microenvironment under confocal microscopy. *PA*, pulmonary artery. *LA*, left atrium. Sketch was made using Biorender.

#### SUPPLEMENTARY FIGURE 9

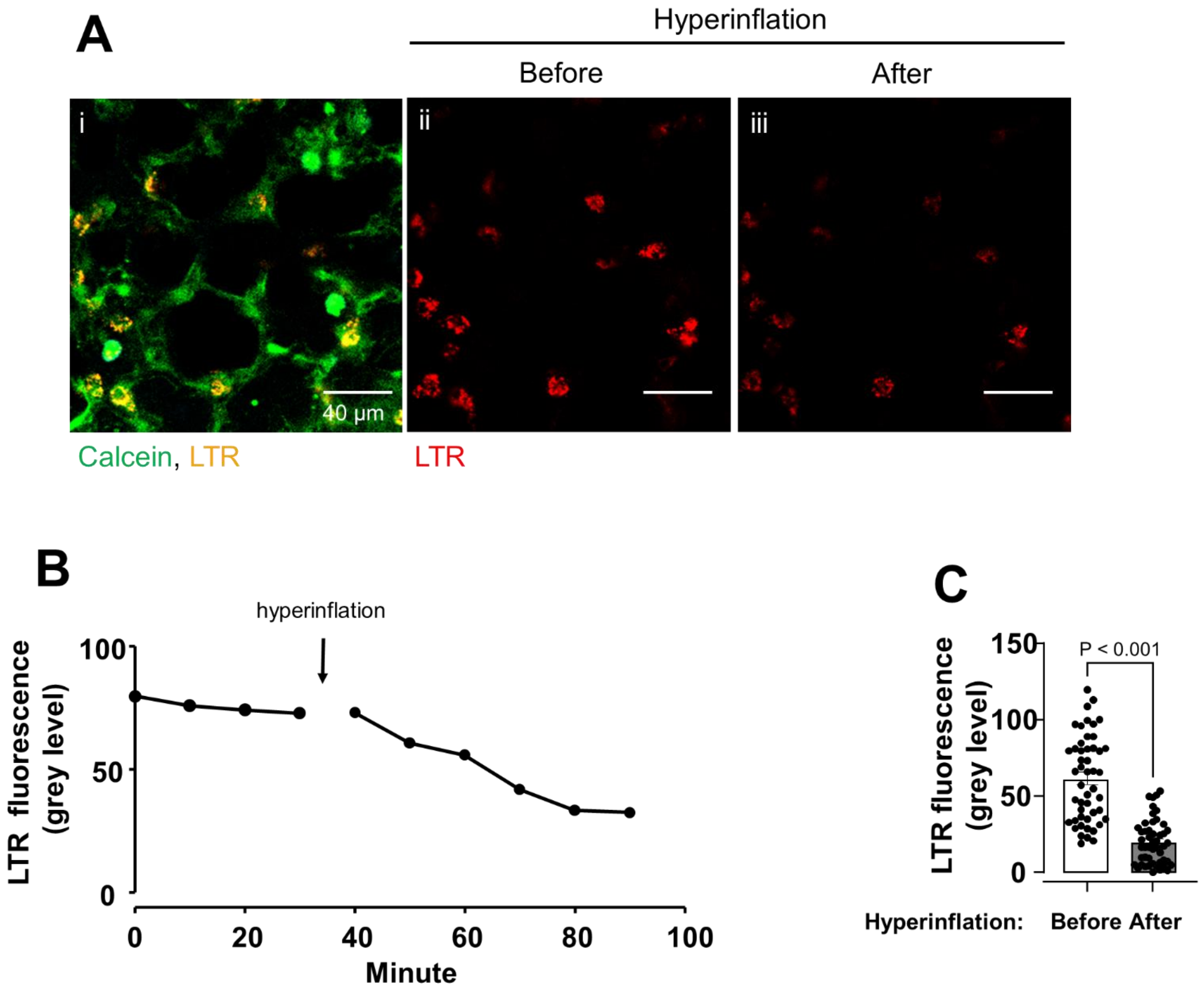


**Supplementary Figure 9. Hyperinflation-induced secretion of surfactant**

**a** Confocal image shows epithelial fluorescence in alveolar epithelium infused with calcein and lysotracker red (LTR) (*i*). Confocal images (*ii-iii*) show LTR fluorescence before and 60 min after hyperinflation.

**b** Tracings from a single LTR+ cell show LTR fluorescence before and after hyperinflation at indicated times.

**c** Bars show paired quantifications of LTR fluorescence (dots) before and 60 min after hyperinflation. Mean±sem, n = 4 lungs, 49 alveolar epithelial type 2 cells**.**

**SUPPLEMENTARY FIGURE 10**

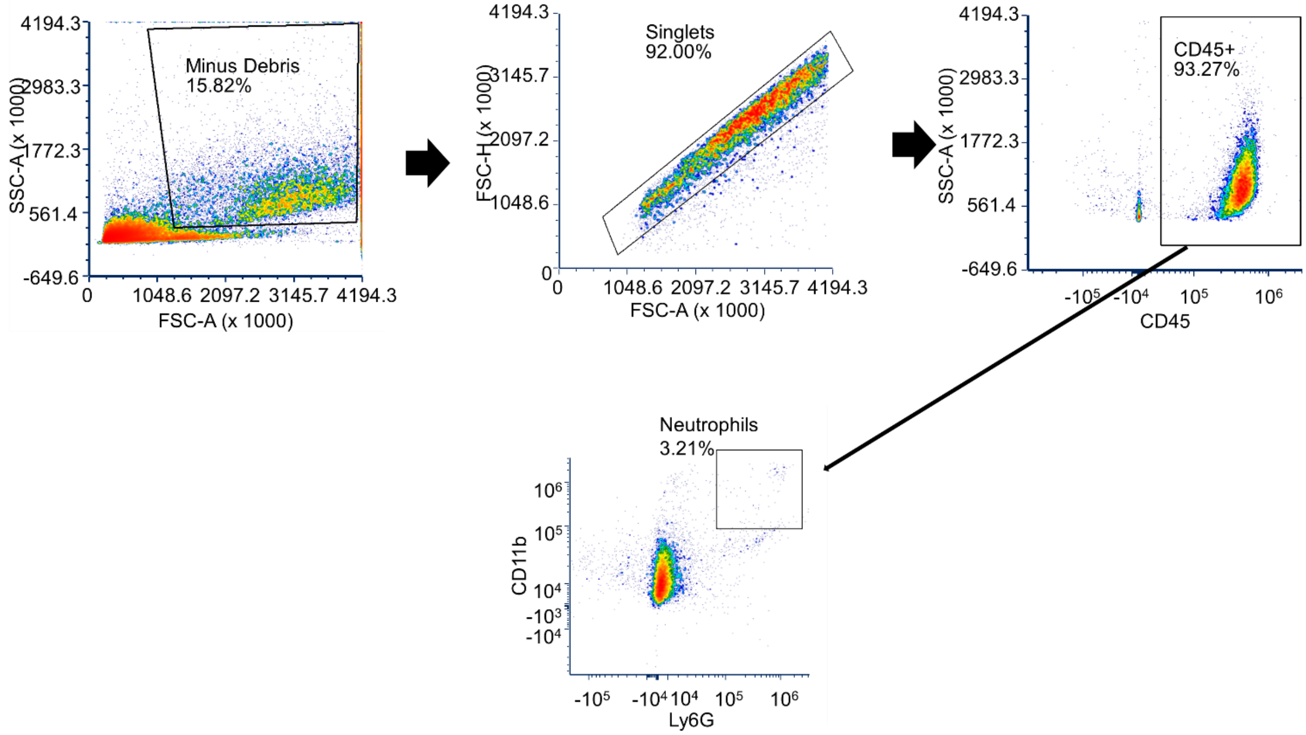


**Supplementary Figure 10. Flow cytometry gating strategy to identify neutrophils in BAL**

**a** Gating strategy used to identify neutrophils from mouse bronchoalveolar fluid (BAL). Mouse was mechanically ventilated at 6 ml/kg for 2 hours then cells were isolated from BAL. After the exclusion of debris and doublets, leukocytes were identified by CD45 staining. A gating strategy was used to identify neutrophils by expression of CD11b-PE-Cy5 and Ly6G-PE-Cy7.

**SUPPLEMENTARY FIGURE 11**

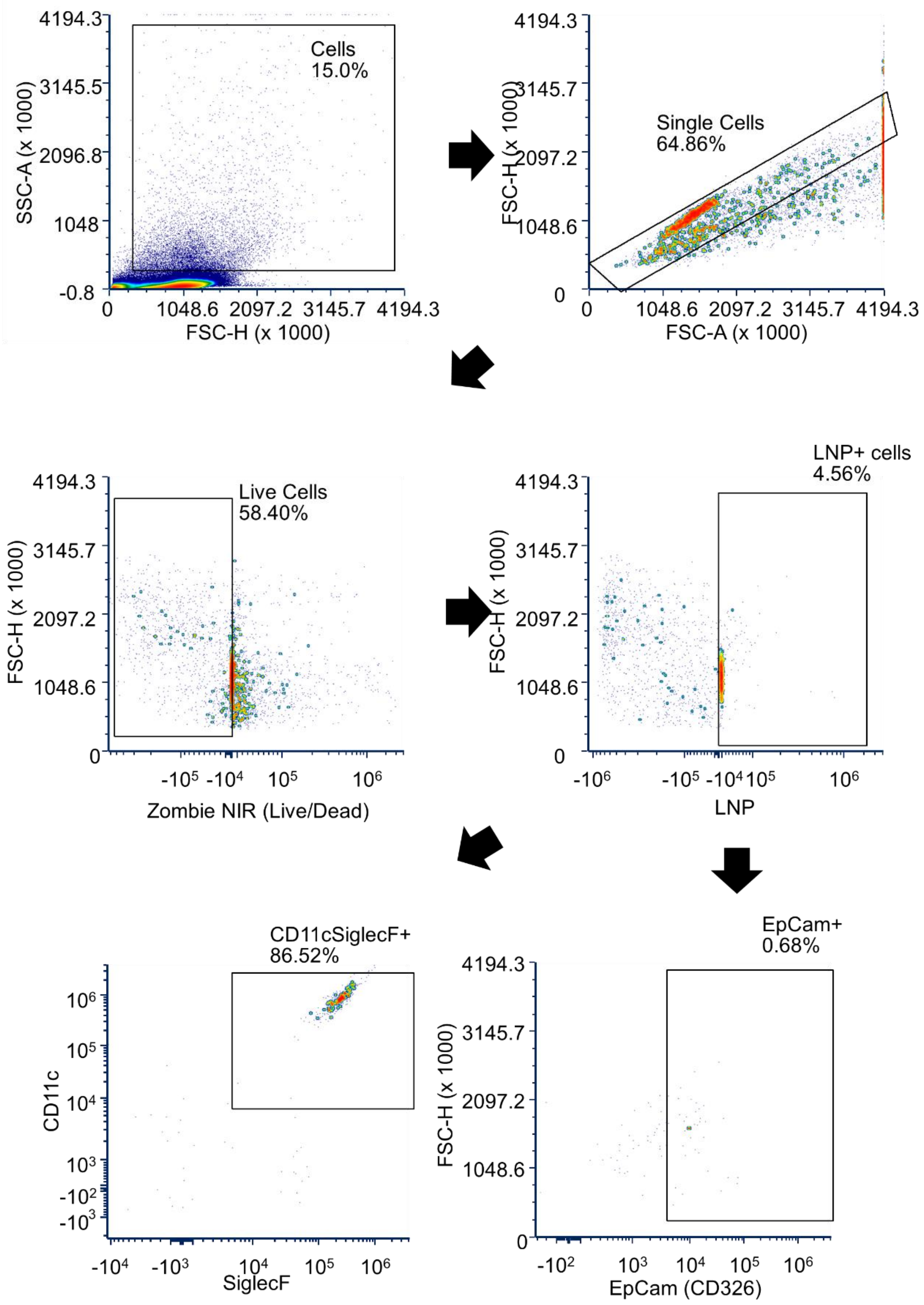

**Supplementary Figure 11. Flow cytometry gating strategy to identify alveolar macrophages and alveolar epithelium in lung homogenates**

Gating strategy used to identify sessile AMs and AE from mouse lung tissue homogenates**.** A mouse was intranasally instilled with rhodamine b dextran 70 kDa LNPs (LNP-FD). Two hours later, cells were isolated from lung homogenates and after the exclusion of debris and doublets. Cells positive for LNP-FD were identified. CD11c-APC and SiglecF-488 and EpCam-PerCP-Cy5.5 were used to identify alveolar macrophages and alveolar epithelium respectively.

#### SUPPLEMENTARY TABLE 1

| Name, Fluorescent Dye | Supplier | Catalogue No. | Stock Conc. | Final Conc. | Lot No. |
|---|---|---|---|---|---|
| CD45, Pacific Blue | Biolegend | 103126 | 0.5 mg/ml | 5 µg/ml | B314223 |
| Ly-6G, PE-Cy7 | Biolegend | 127618 | 0.2 mg/ml | 2 µg/ml | B351626 |
| CD11b, PE-Cy5 | Biolegend | 101210 | 0.2 mg/ml | 2 µg/ml | B260953 |
| CD11c, APC | Biolegend | 117310 | 0.2 mg/ml | 2 µg/ml | B278343 |
| SiglecF, PE | Biolegend | 155506 | 0.2 mg/ml | 2 µg/ml | B301117l |
| Siglec1 (CD169) | Thermo Fisher | PA5-84155 | 0.17 mg/ml | 3.4 µg/ml | ZA4189480 |
| CD203a, Alexa 647 | BioRad | MCA1973A647 | 0.05 mg/ml | 5 µg/ml | 152566 |
| Goat anti-rabbit IgG, Alexa Fluor 488) | Molecular Probes | A-11008 | 1 mg/ml | 100 µg/ml | 2743033 |
| SiglecF, Alexa Fluor 488 | Invitrogen | 53-1702-80 | 0.2 mg/ml | 2 µg/ml | 2072850 |
| EpCam, PerCP-Cy5.5 | Biologend | 118219 | 0.2 mg/ml | 2 µg/ml | B406473 |
| TNFR1, Alexa Fluor 633 | AbD Serotec | MCA2350 | 10 mg/ml | 40 µg/ml | n/a |

**Supplementary Table 1. Table of antibodies used in imaging and flow cytometry studies**

Final concentration (conc.) indicates concentration used in experiments.

#### SUPPLEMENTARY TABLE 2

| Donor Network | Donor | Age | Gender | Smoking | Substance use | Ethnicity |
|---|---|---|---|---|---|---|
| LiveOnNY | D536 | 34 | F | Y | Marijuana, Amphetamines, PCP, Cocaine, and Barbituates | Hispanic/Latino |
| LiveOnNY | D546 | 52 | M | Y | THC, Cocaine, Opiates | Hispanic/Latino |
| LiveOnNY | D567 | 46 | M | Y | Cocaine, EtOH | Hispanic/Latino |
| LiveOnNY | D672 | 82 | M | Y | EtOH | Hispanic/Latino |

**Supplementary Table 2. Participants**

Demographic data for each donor is indicated.
